# Quadrupia: Derivation of G-quadruplexes for organismal genomes across the tree of life

**DOI:** 10.1101/2024.07.09.602008

**Authors:** Nikol Chantzi, Akshatha Nayak, Fotis A. Baltoumas, Eleni Aplakidou, Shiau Wei Liew, Jesslyn Elvaretta Galuh, Michail Patsakis, Camille Moeckel, Ioannis Mouratidis, Saiful Arefeen Sazed, Wilfried Guiblet, Austin Montgomery, Panagiotis Karmiris-Obratański, Guliang Wang, Apostolos Zaravinos, Karen M. Vasquez, Chun Kit Kwok, Georgios A. Pavlopoulos, Ilias Georgakopoulos-Soares

## Abstract

G-quadruplex DNA structures exhibit a profound influence on essential biological processes, including transcription, replication, telomere maintenance, and genomic stability. These structures have demonstrably shaped organismal evolution. However, a comprehensive, organism-wide G-quadruplex map encompassing the diversity of life has remained elusive. Here, we introduce Quadrupia, the most extensive and well-characterized G-quadruplex database to date, facilitating the exploration of G-quadruplex structures across the evolutionary spectrum. Quadrupia has identified G-quadruplex sequences in 108,449 reference genomes, with a total of 140,181,277 G-quadruplexes. The database also hosts a collection of 319,784 G-quadruplex clusters of 20 or more members, annotated by taxonomic distributions, multiple sequence alignments, profile Hidden Markov Models and cross-references to G-quadruplex 3D structures. Examination of G-quadruplexes across functional genomic elements in different taxa indicates preferential orientation and positioning, with significant differences between individual taxonomic groups. For example, we find that G-quadruplexes in bacteria with a single replication origin display profound preference for the leading orientation. Finally, we experimentally validate the most frequently observed G-quadruplexes using CD-spectroscopy, UV melting, and fluorescent-based approaches. Quadrupia is publicly available through https://www.pavlopoulos-lab.org/quadrupia.

## Introduction

DNA G-quadruplexes (G4s) are one of the most thoroughly studied non-B DNA structure. G4s are commonly found in GC-rich areas of a genome and are characterized by Hoogsteen base pairs, in which hydrogen bonds link four guanine bases into a square planar formation known as a G-quartet. Multiple G-quartets, stacked on top of each other, lead to the formation of G4 structures (1) (**Figure 1a**). Their presence and formation in telomeric regions provided early evidence that G4s are a native DNA conformation (2). Furthermore, an array of studies have shown that G4s are involved in processes such as gene regulation (3–13), alternative splicing modulation (14–16), 3D genome structure organization (17–19), and translation (20–25), among other functions. Additionally, G4s are linked to genomic instability and have been associated with several diseases, including cancer and neurodegenerative disorders (26–31).

**Figure 1:**
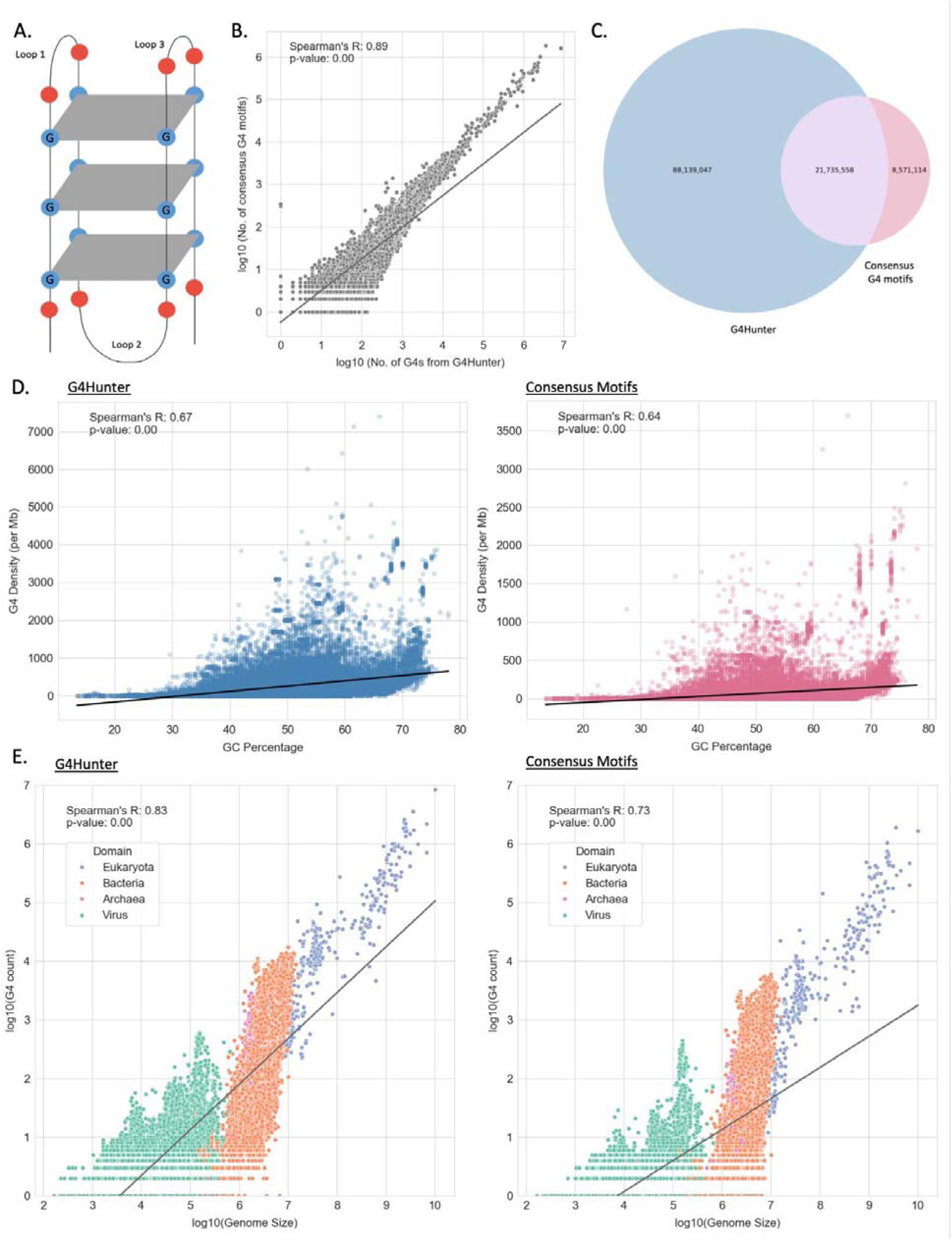
Characterization of G4s across 108,534 genomes. **A)** Schematic illustration of a G4. **B)** Scatter plot displaying the association in the number of G4s detected per species with each of the two methods. **C)** Venn diagram showing the number of shared G4s by the two methods. **D)** Association between GC content per genome and number of G4s observed per Million base pairs for results from G4Hunter (left) and consensus G4 motifs (right). **E)** Association between genome size and number of G4s detected for results from G4Hunter (left) and consensus G4 motifs (right). Every dot represents a genome and coloring reflects the taxonomic subdivision in the three domains of life and viruses.

There are several established methods for detecting G4s *in vitro* and *in vivo* and characterizing their formation kinetics and stability (32). *In vitro* approaches include chemical and biophysical methods such as nuclear magnetic resonance (NMR) (33, 34), circular dichroism (35, 36) and UV thermal melting (37). G-quadruplex sequencing methods (G4-seq and rG4-seq) use the property of stable G4 structures to impede DNA and RNA polymerase progression *in vitro*, enabling genome-wide or transcriptome-wide high-resolution detection of putative DNA and RNA G4s through high-throughput sequencing (38–41). To detect G4s *in vivo*, immunostaining with antibodies specifically designed against G4s (Biffi et al. 2013) and ChIP-seq genome-wide approaches have been implemented (9). Imaging of G4 formation has been demonstrated in live cells using fluorescence imaging (42, 43). Such experimental approaches have enabled the derivation of patterns enabling the prediction of sequences that can adopt G4 DNA structures.

Early computational studies showed that conserved sequence motifs can accurately capture a significant proportion of potential G4-forming regions (44). Since then, the consensus G4 motif, G≥3N1–7G≥3N1–7G≥3N1–7G≥3, has been used to quantify the number and distribution of G-quadruplexes and is utilized in numerous available methods and studies (Kikin et al. 2006; Huppert and Balasubramanian 2007; Zhang et al. 2008; Wong et al. 2010; Cer et al. 2013). However, it has become evident that certain G4s do not conform to this consensus motif. Thus, additional computational approaches have since been implemented to capture a broader range of putative G4-forming sequences, including G4Hunter (45–47).

Previous work has characterized the distribution and frequency of G4s in species from multiple taxonomies. In eukaryotes, G4s are enriched at *cis*-regulatory elements (3–7), including in promoters, enhancers, and CTCF binding sites, while in higher eukaryotes they have also emerged in proximity to splice sites to modulate alternative splicing (14). In bacteria, an analysis of 1,627 genomes revealed significant differences in the frequencies of G4s, with Deinococcota displaying the highest density in their genome (48), while another study revealed enrichment of G4s in functional elements when examining eighteen prokaryotic genomes (49). When investigating the genomes of archaea and viral species large differences in the frequencies and functions of G4s were also reported (50–53).

To date, there are a few databases available incorporating the identification of potential G4 DNA-forming sequences in different organisms; however, all available datasets cover a small number of species. Zhong et al. (2023) constructed G4Bank, a G4 database containing six million G4s across thirteen species (54), while GAIA currently harbors G4s for 61 different organismal genomes (55). Another database named G4Atlas contains only 238 experimental G4s from 10 species (56). Other databases are focused only on a small set of species or are providing a set of experimentally validated G4s (57–61), yet none of these databases contain a wide range of genomes, representing species across the taxonomic subdivisions in the tree of life. Thus, there is a gap in extensive evolutionary research on G4s as there is a lack of genome-wide maps of predicted G4s across all species with a reference genome available.

Here, we present “Quadrupia”, the largest and, to our knowledge, most comprehensive database of putative G4 DNA-forming sequences to date, covering 108,534 species and encompassing 87,160,084 putative G4 DNA-forming sequences. The experimental validation of highly prevalent G4s across the tree of life suggests that our predictions in the database are accurate. We identify clusters of G4 DNA-forming sequences based on sequence identity and characterize the preference of each cluster in different taxonomies. We also perform G4 DNA analyses across the taxonomic subdivisions and at individual species, observing marked differences in G4 DNA density and distribution across functional genomic elements. For example, we discovered that G4s are strongly enriched in the leading strand relative to the lagging strand orientation in bacterial genomes and are enriched in a subset of splice junctions in specific phyla. We expect that our work will open up opportunities for researchers to delve into the functional, regulatory, and evolutionary origin of G4 DNA, using the plethora of available complete organismal genomes.

## Methods

### Data retrieval and parsing

Complete genomes were downloaded from the GenBank and RefSeq databases (62, 63). A total of 108,449 complete genomes were analyzed and integrated in the database. For each genome, the associated files for RNA and coding regions as well as the GFF gene annotation file were downloaded. Replication origins for bacteria were derived from Doric Database for complete assemblies, for circular topology and for single origin of replication types (64).

### Identification of potential G4 DNA-forming sequences

Regular expressions were employed to generate genome-wide G4 maps, as well as for RNA and coding regions. For each species’s genome, RNA, and coding regions, we generated the genome-wide DNA G4 maps using a regular expression of the consensus G4 motif (G ≥ 3N1-7G ≥ 3N1-7G ≥ 3N1-7G ≥ 3) (65). G4s were also detected using G4hunter with parameters of a window -w =25 and -s 1.5 (46, 66) and processed by a custom Python script to transform the data into a processable tabular format. Since G4s were extracted from both GenBank and RefSeq assemblies resulting in some duplicates, unique G4s were obtained by prioritizing RefSeq assemblies over GenBank due to a more comprehensive annotation of the RefSeq assemblies. G4s with overlapping coordinates were merged into a single sequence before performing further analysis. Shared G4s between the G4Hunter and the regular expression methods were estimated using a minimum overlap of 50% between motifs detected from the two methods. The requirement for a minimum of 50% overlap needs to be satisfied by any one of the two sequences for them to be considered common between the two methods.

### Estimation of G4 density across genomes and genomic subcompartments

G4 density was calculated as the number of G4 bps over the number of base pairs examined. The average G4 density was calculated between organismal genomes within taxonomic groups. The G4 density was also examined across genomic subcompartments and was calculated as the length of overlap of G4s with each of the subcompartments divided by the total length of the subcompartments. The subcompartment coordinates were obtained from the corresponding GFF files and overlapping annotations within a subcompartment were merged. The mean G4 density was calculated across species belonging to the same taxonomy either across the genome or in genomic subcompartments.Assembly acccessions associated with a GFF file were included in this analysis. In species that were missing RNA features, due to redundancy, the missing genomic subcompartments were added using a custom python script. Species that did not have any relevant genomic compartment annotated in their GFF files were excluded from the analysis. The mean G4 density was log-transformed for the categorical plots and the heatmaps in Fig 3. To reduce noise, phyla with less than five species were excluded from the heatmap (**Fig 3B**).

### Estimation of G4 density relative to TSSs, TESs, replication origins and splice sites

To investigate the relationship between G4 sites and TSSs or TESs, we generated local windows of 500bps long around TSSs/TESs and measured the distribution of G4 bps across the window. To that end we use the gene coordinates, extracted from the corresponding GFF files. The enrichment was calculated as the sum of G4 occurrences at each relative position across organismal genomes over the mean of the resulting number of occurrences across the generated window. Confidence intervals were calculated as the 2.5% lowest and 97.5% highest percentile from Monte-Carlo simulations with replacement (N=1,000), in which we randomly picked an equal number of species from the domain, kingdom or phylum that was studied.

### Origin of Replication

We investigated the distribution of G4s across bacterial genomes with circular chromosomes. To that end, we downloaded the origin of replication from the single-partite datasets from the Doric database. Each organismal genome in Doric was annotated with the start and end of origin of replication. For each available chromosome, we derived the halfway distance from the oriC as (start+end)/2. Afterwards, we binned the chromosomal sequence in 1001 bins, with bin 500 corresponding to oriC. Due to the circular nature of the chromosome, we used the distance metric d(P,Q)=min(|P-Q|, ChromosomeSize-|P-Q|), in which P, and Q are any two genomic positions. Consequently, the bin 1001 coincides with - is identical to bin 0, whereas 500 bin is identical to the replication terminus. We associated each G4 sequence to a specific bin (in rare cases a sufficiently large G4 was assigned to more than 1 bin), by calculating the circular distance on the binned chromosome for that particular G4 from oriC. The resulting binned distribution, consisted of bins containing the total number of G4s that have a distance of d units from oriC. Note that looking at the generated figure, a positive distance d>0 would translate to G4 sequence being located on the left rather than on the right half-circle; since oriC is always placed at pi/2, assuming counterclockwise orientation. Finally, we divide each position by the total mean to estimate the local enrichment of G4s. This process was repeated separately for G4s located on forward and on reverse strand, respectively, as well as for both G4Hunter and regex methodologies. These distributions were used to generate the G4 polar scatterplots in relation to origin of replication, using matplotlib polar coordinate system. Furthermore, we extracted the GC-Skew for each organismal genome. To achieve this, we split the chromosome into windows of 25bp and calculated the GC-Skew using the formula G-C/G+C. Then, following the process above, we assigned to each bin of the circular chromosome the average GC-Skew of all the windows belonging to a particular bin.

### Biophysical Properties

We investigated the biophysical properties of G4 motifs derived from the regex algorithm and G4Hunter, by decomposing each G4 into consecutive G-runs. A G-run is defined as any subsequence of G4 containing at least more than 3 consecutive guanines. Any G4 is composed of consecutive G-runs interrupted by a sequence that violates the pattern. We defined the loop as the union of all the nucleotide sequences between the G-runs. We used regular expressions to extract the G-runs and intervening loops for each G4 motif and thereafter we calculated the total frequency of each G-run length as well as the frequencies of the various loop lengths associated with each G4. Analyses were performed to compare the overall distribution across taxonomies, and at individual taxonomies.

### Circular dichroism (CD) spectroscopy

Reaction samples of 2 ml containing 5 μM oligos were prepared in 10 mM LiCac (pH 7.0) and 150 mM KCl or LiCl. The samples were mixed well and denatured by heating them at 95°C for 5 minutes, and then cooled down to room temperature for renaturation. Data measurements were obtained using a Jasco CD J1500 spectrometer and a quartz cuvette with a path-length of 1 cm. The samples were scanned at 2 nm intervals starting from 220 nm and ending at 310 nm. The resulting data were blanked and normalized to obtain the mean residue ellipticity. The samples were scanned 3 times and data were averaged. Analysis of all data was done in Spectra Manager™ Suite and Microsoft Excel.

### UV melting spectroscopy

Reaction samples were prepared and renatured in accordance with the CD experiment above for the KCl condition, and the UV melting assay was carried out in a Cary 3500 UV-Vis Multicell Peltier spectrometer and a 1 cm path-length quartz cuvette sealed with thread seal tape. Data measurements were acquired every 0.5°C increments from 20 °C to 95 °C at 295 nm. All recorded data were blanked and smoothed by averaging every 10 °C. Data analysis was performed in Microsoft Excel.

### G4 ligand-enhanced fluorescence spectroscopy

Reaction samples of 1 μM oligos were set up in 10 mM LiCac (pH 7.0) and 150 mM KCl or LiCl to a total volume of 100 μl and mixed well. Renaturation was conducted by heating the samples at 95°C for 5 minutes and cooling down at room temperature for 15 minutes. After the addition of 1 μM of NMM or ISCH-OA1 ligand, the fluorescence spectra of the samples were collected using a HORIBA FluoroMax-4 fluorescence spectrophotometer and a 1 cm path-length quartz cuvette. The excitation wavelengths for NMM and ISCH-OA1 were set to 394 and 570 nm respectively, and the spectra were collected at 550-750 nm for NMM and 590-750 nm for ISCH-OA1. Entrance slit was set to 5 nm and exit slit was set to 2 nm, and the spectra were collected every 2 nm and smoothed every 3 data points. All recorded data were analyzed using Microsoft Excel.

### Sequence clustering

Sequence clustering was performed for the combined datasets of regex- and G4hunter-derived G4 sequences. Any coordinate overlaps between the two datasets were detected as described above, and for each set of overlapping sequences, the longest was chosen as the representative candidate. Clustering was performed using the Linclust algorithm (67) implemented in MMseqs2 (68). Linclust was chosen as it is a kmer - based method capable of clustering hundreds of millions of sequences in linear time regardless of kmer length, thus being the most efficient solution for G4s. Clustering was performed using an 80% sequence identity cutoff and a 90% bidirectional alignment coverage threshold, while the minimum cluster size was set at 20 members. These cutoff values were chosen to reflect the conservative nature of G4 motifs and, at the same time, optimize the number of clustered versus unclustered (singleton) sequences. Each G4 cluster was used to produce a multiple sequence alignment (MSA) with MAFFT (69), applying the directional adjustment parameter to consider reverse complementarity and ensure proper bidirectional alignment. The centroid sequences of each cluster, as generated by MMseqs2, were used to guide MSA generation and were defined as the representative sequences of the clusters. The MSAs were used to generate profile Hidden Markov Models (pHMMs) with HMMER v. 3.3.2 (70).

### Identification of structural representatives

A dataset of experimentally determined G-quadruplex 3D structures was constructed by parsing the records of the OnQuadro database (61) (retrieved in May 06, 2024) and matching them to their corresponding PDB (71) entries. The DNA sequences of this dataset were searched against the representative sequences of the G4 clusters using BLAST+ (72). Sequence hits were identified using a sequence identity cutoff of 80% and an alignment coverage threshold of 90% with respect to the shortest sequence. The structure with the top sequence hit to a cluster (largest sequence identity, followed by largest alignment coverage), was chosen as the cluster’s structural representative.

### Database Implementation

The front end of Quadrupia was implemented in HTML, CSS, and JavaScript. The back end is supported by the Apache web server and the Slim Framework v. 4.0, with server-side operations handled by PHP and, when required, Python. The genome metadata is stored in a MySQL relational database. The Quadrupia website layout is designed using the Bootstrap v. 5 framework, jQuery, and the DataTables library. Data visualization is performed using MSAviewer (73) for MSAs, SkyLign (74) for sequence logos and Molstar (75) for 3D structures. Sequence queries are performed using BLAST+ (72) for sequences and *nhmmscan* (76) for pHMMs. An additional option to perform motif-based queries is also offered, implemented using Python.

## Results

### Identification of potential G4-forming sequences in organismal genomes across the tree of life

We examined 108,459 organismal reference genomes spanning the three domains of life and viruses and identified putative G4 DNA-forming sequences in each of them using two methods, namely the consensus G4 motif and the predicted G4s from the state-of-the-art G4 detection method G4-hunter (46). We find an average of 279.43 G4s per genome with the consensus G4 motif method and 1,013.05 per genome with the G4hunter-based method, observing a high degree of concordance between the number of G4s detected with each method for each genome (Spearman correlation r = 0.89, p-value = 0, **Figure 1b-c; Supplementary Figure 1a**). Consensus G4 sequences and G4 sequences found using the G4hunter method that have at least a 50% overlap were considered common G4 sequences found using both methods (**Figure 1c**).

G4s are highly GC-rich sequences; we therefore investigated the association between each organismal genome’s GC content and the number of G4s detected. We observe a strong correlation between the density of G4s and the GC content of each species, with both G4 detection approaches (**Figure 1d**, Spearman correlations r=0.67; p-value=0 for results from G4Hunter and r=0.64; p-value=0 for consensus G4 motif method). We also find a large disparity in the G4 density between genomes, for a given GC content, indicating that GC content can only partially account for the G4 density differences between organismal genomes (**Figure 1d**).

Next, we investigated the number of G4s observed across the taxonomies as a function of genome size. As expected, larger genomes harbored a larger number of G4s, with eukaryotes and viruses having the highest and lowest number of G4s, respectively (**Figure 1e**). We also find that there is a large dispersion within the domains of life and viruses (**Figure 1e**). We report that the species with the highest G4 density in their genomes based on both approaches are Grapevine fleck virus and Alcea yellow mosaic virus, with densities of 104.57 and 93.27 G4s per kB, respectively, based on the consensus G4 motif approach, and densities of 438.92 and 371.51 G4s per kB, respectively, based on G4Hunter. We also examined the association between the proportion of the genome covered by genes and the genomic density of G4s per genome and observed a weak negative correlation in the case of eukaryotes. (**Supplementary Figure 1b** Spearman correlations r=-0.25; p-value=0 for results from G4Hunter and r=-0.34; p-value=0 for consensus G4 motif method).

### Characterization of G4s across taxonomic subdivisions

Next, we investigated differences in the distribution and density of G4s between taxonomic subgroups. We first examined the association between the genome size and the density of G4s for organisms in the three domains of life and viruses. Interestingly, we find that there is a moderately positive correlation between the density of G4s and the genome size in bacteria (Spearman correlations r=0.44; p-value=0 for results from G4Hunter and r=0.51; p-value=0 for consensus G4 motif method) and eukaryotes (Spearman correlations r=0.41; p-value=0 for results from G4Hunter and r=0.52; p-value=0 for consensus G4 motif method). In archaea, results from G4Hunter show no correlation while results from the consensus G4 motif method show a very weak positive correlation (r=0.25; p-value=0). In viruses, no correlation between the density of G4s and the genome size is observed in results from either method (**Figure 2a**). We also examined the G4 density in the three domains of life and viruses. We report that the highest and lowest density of G4s is observed in eukaryotes (327.03 G4s per Mb) and bacteria (147.81 G4s per Mb), respectively, based on the results from G4Hunter. However, based on the consensus G4 motifs, the highest and lowest density of G4s is observed in eukaryotes (82.76 G4s per Mb) and archaea (26.40 G4s per Mb) (**Figure 2b**).

**Figure 2:**
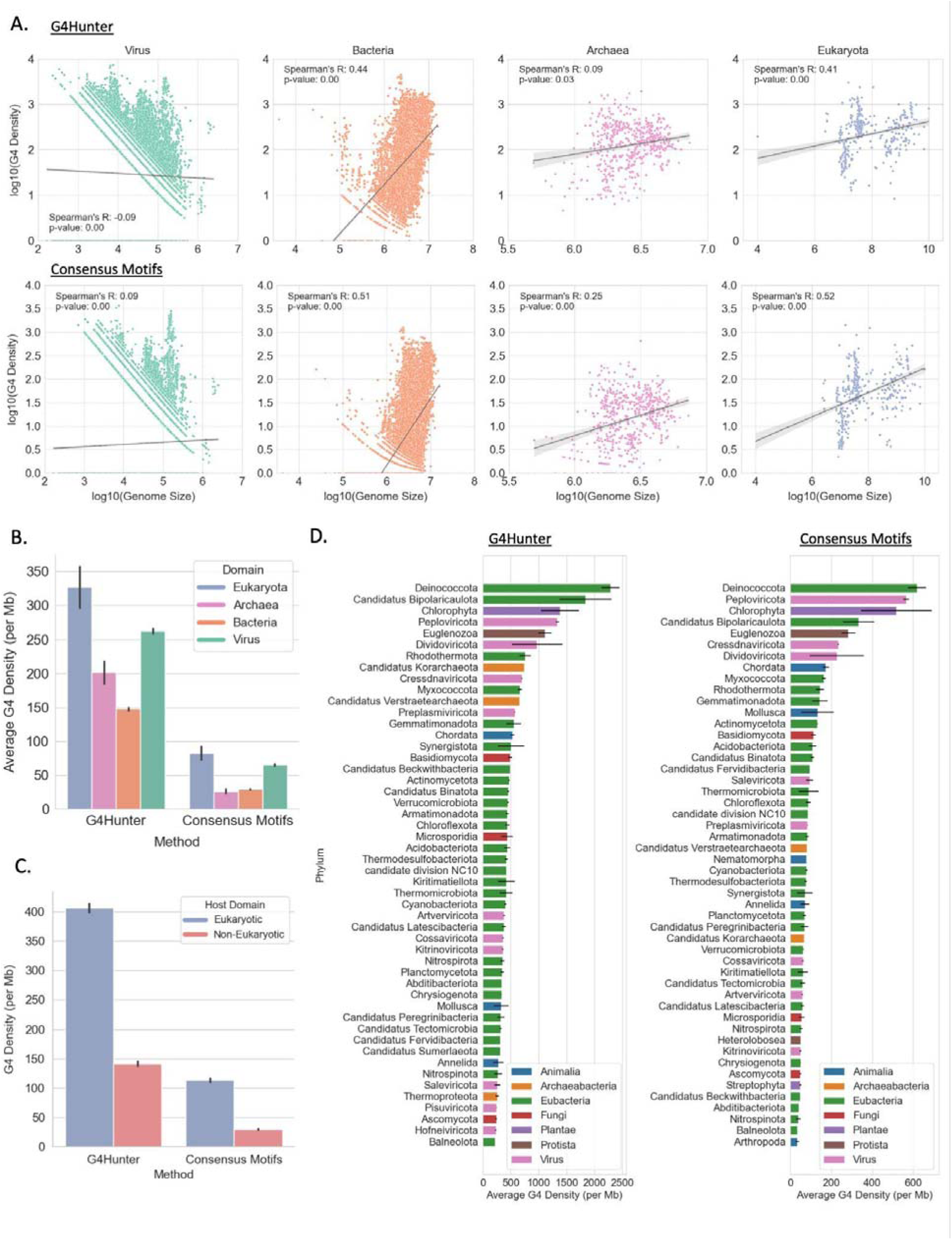
Taxonomic characterization of G-quadruplexes across the tree of life. **A)** Association between genome size and number of G4s detected, stratified by taxonomic subdivision in the three domains of life and viruses. **B)** Density of G4s in each domain of life and viruses. **C)** Association between G4 densities of viral genomes and their host domains**. D)** Density of G4s in each phylum. Error bars show standard error.

We further subdivided this analysis at the phylum level. The average G4 density varied between phyla belonging to each domain, with about 2,285-fold variation in bacteria, 1,382-fold in eukaryotes, 1,336-fold in viruses, and 742-fold in archaea. We observe that the top five phyla with the highest G4 density are Deinococcota, Candidatus Bipolaricaulota, Chlorophyta, Peploviricota, and Euglenozoa in both G4Hunter and the consensus G4 motif approaches (**Figure 2d**), indicating strong concordance between the two G4 detection approaches used. Additionally, the phyla with the highest G4 density belong to different domains of life and viruses and to different kingdoms (**Figure 2d; Supplementary Figure 3**), highlighting the significant variability in G4 density between phyla belonging to the same taxonomic supergroup. These findings underscore the profound differences in G4 density in organismal genomes across the tree of life.

Previous studies have shown a correlation between G-quadruplex sequence composition in viruses and their hosts (77). We therefore investigated if the viruses with the highest G4 densities also have hosts with a high G4 density. Out of the 63,658 viral genomes that we analyzed in this study, we selected the top 100 viral genomes with the highest G4 density. The majority of the viruses were found to be part of the phylum Peploviricota (73% in G4Hunter results and 94% in results obtained from consensus G4 motifs) having vertebrate hosts, followed by phylum Kitrinoviricota (11% in G4Hunter results and 4% in results obtained from consensus G4 motifs) having plant and fungal hosts. Despite Peploviricota and Kitrinoviricota accounting for only 2.2% and 2.8%, respectively, of all the viral genomes we studied, they collectively constituted a significant proportion of the top 100 viruses with the highest G4 density. Notably, both have eukaryotic hosts which we found to have the highest G4 density among the three domains of life **(Figure 2b)**. Further investigating the G4 densities of all viral genomes with hosts across the three domains of life, we found viruses with eukaryotic hosts exhibited the highest average G4 density **(Figure 2c; Supplementary Table 1)**, followed by archaeal and bacterial hosts **(Supplementary Figure 1c)**. A two-sample t-test indicated a statistically significant difference (p-value=0.0 for G4Hunter results and p-value=0.0 for consensus G4 motifs) in the average G4 densities between the viruses with eukaryotic and prokaryotic hosts, with the viruses with eukaryotic hosts having substantially higher G4 densities. These results indicate a possible association between the G4 densities of viruses and their hosts.

### Genomic distribution of G4s in genomic subcompartments across organismal genomes

G4s are unevenly distributed in the human genome, and are enriched in specific sub-compartments in which they have functional roles associated with transcription and translation (3–12, 14, 15, 78, 79). We therefore investigated the extent to which the G4 density varied within organismal genomes, in different genomic sub-compartments, and if such differences were influenced by the taxonomic group studied.

To that end, we examined the G4 density across the genome and in genic, exonic, and coding regions for each organismal genome across the different taxonomic groups. For the three domains of life and viruses, we observed that the density of G4s is highest when examining the whole genome, rather than any specific genomic subcompartment (**Figure 3a**). The results were consistent across both methodologies based on G4Hunter results and the consensus motifs. Similarly, when examining different phyla based on the results from G4Hunter, genomic-wide G4 distribution has the highest average G4 density per Mb. However, there are some notable exceptions. In particular, the eukaryotic phylum Chordata indicates a higher G4 density in exonic and coding sequences (CDS) (**Figure 3b**). Furthermore, several viral phyla, such as Dididoviricota, display a higher genic G4 density than their genomic counterparts (**Figure 3b**). However, this analysis did not examine the G4 positioning relative to functional genomic sites. Based on G4Hunter methodology, we also find that in eukaryota, the median 24.5% of total genes per species harbor one or more G4s whereas, in archaea, 11.77% of total genes have at least of G4 sequence per species, followed by bacteria with a 7.7% and viruses 4.7% (**Supplementary Figure 2**). Furthermore, if we partition the genes into the two mutually exclusive sets of protein-coding and non-coding, we find in eukaryotic organisms an even higher median overlap of 25%, while across all four domains, the non-coding overlap is lower than its corresponding protein-coding (**Supplementary Figure 2)**. These findings indicate that G4s are highly prevalent in sites associated with transcriptional regulation and transcription.

**Figure 3:**
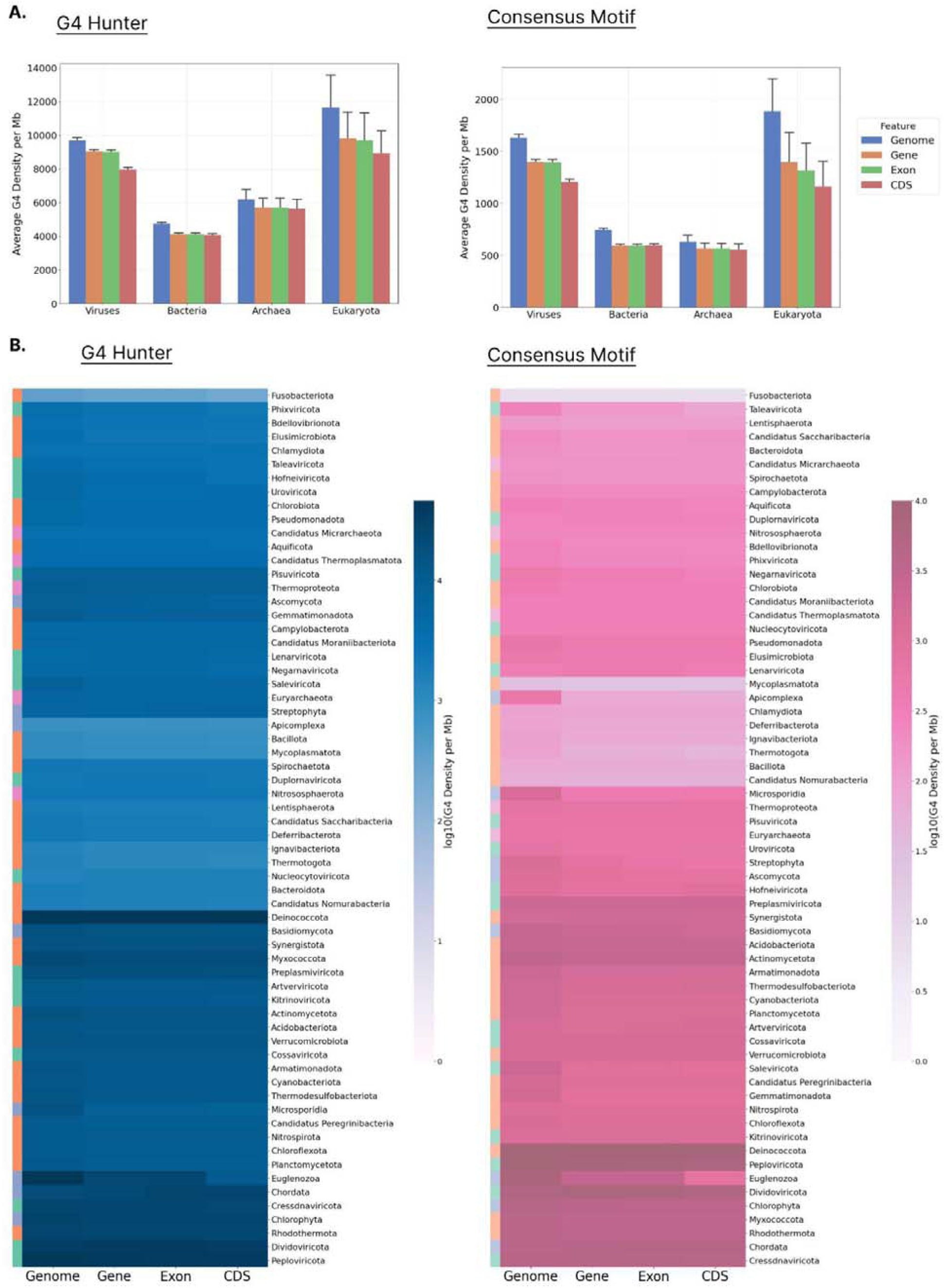
G4 density in different genomic sub-compartments for organisms across the tree of life. **A)** G4 density for the three domains of life and viruses at the genome, genic, exonic and coding regions. **B)** G4 density of organisms belonging to different phyla at the genome, genic, exonic and coding regions.

### Enrichment of G4s relative to transcription start and end sites and splice sites

We next investigated how G4 sequences are distributed concerning transcription start sites (TSSs) and transcription termination sites (TESs). By comparing the varying frequencies of G4s near these sites across different taxonomic groups, we sought to understand potential differences in the regulatory roles of G4s in gene expression among taxa. For G4Hunter-based G4 detection, we observe that there is an enrichment of G4s in the promoter upstream regions for bacteria (1.84-fold enrichment), eukaryotes (1.62-fold enrichment), archaea (1.63-fold enrichment) and viruses (1.54-fold enrichment), while similar results were observed from G4s derived using the regular expression-based algorithm (**Supplementary Figure 4**-5). Relative to the TES, we found strong enrichments using the G4Hunter algorithm for archaea (1.57-fold), bacteria (2.18-fold), and viruses (1.77-fold), with weaker enrichment also observed for eukaryotes (1.18-fold), results that were consistent when using the regular expression-based algorithm (**Supplementary Figure 4**-5).

When examining G4s separately in the template and non-template strands, we found that there were large differences in their distribution in all domains of life, both relative to the TSSs and TESs (**Figure 4b; Supplementary Figure 6**). For instance, G4 sequences found in bacteria and archaea were predominantly located downstream of the TES on the template strand (enrichments of 2.78-fold and 2.34-fold with the G4Hunter algorithm) relative to the non-template strand (enrichments of 1.54-fold and 1.56-fold), while G4s on the non-template strand were more commonly found in regions preceding the TES (Kolmogorov-Smirnov test, p-values<0.0001, Bonferonni corrected p-values) (**Figure 4b**). Interestingly, G4s were depleted in the template strand in the vicinity of the TES, but not in the non-template strand (**Figure 4b**). We further separated organisms from the three domains of life and viruses into the different phyla. We observe that the distribution of G4s relative to TSSs and TESs are highly variable between the different phyla (**Figure 4b-c; Supplementary Figure 7**-8).

**Figure 4:**
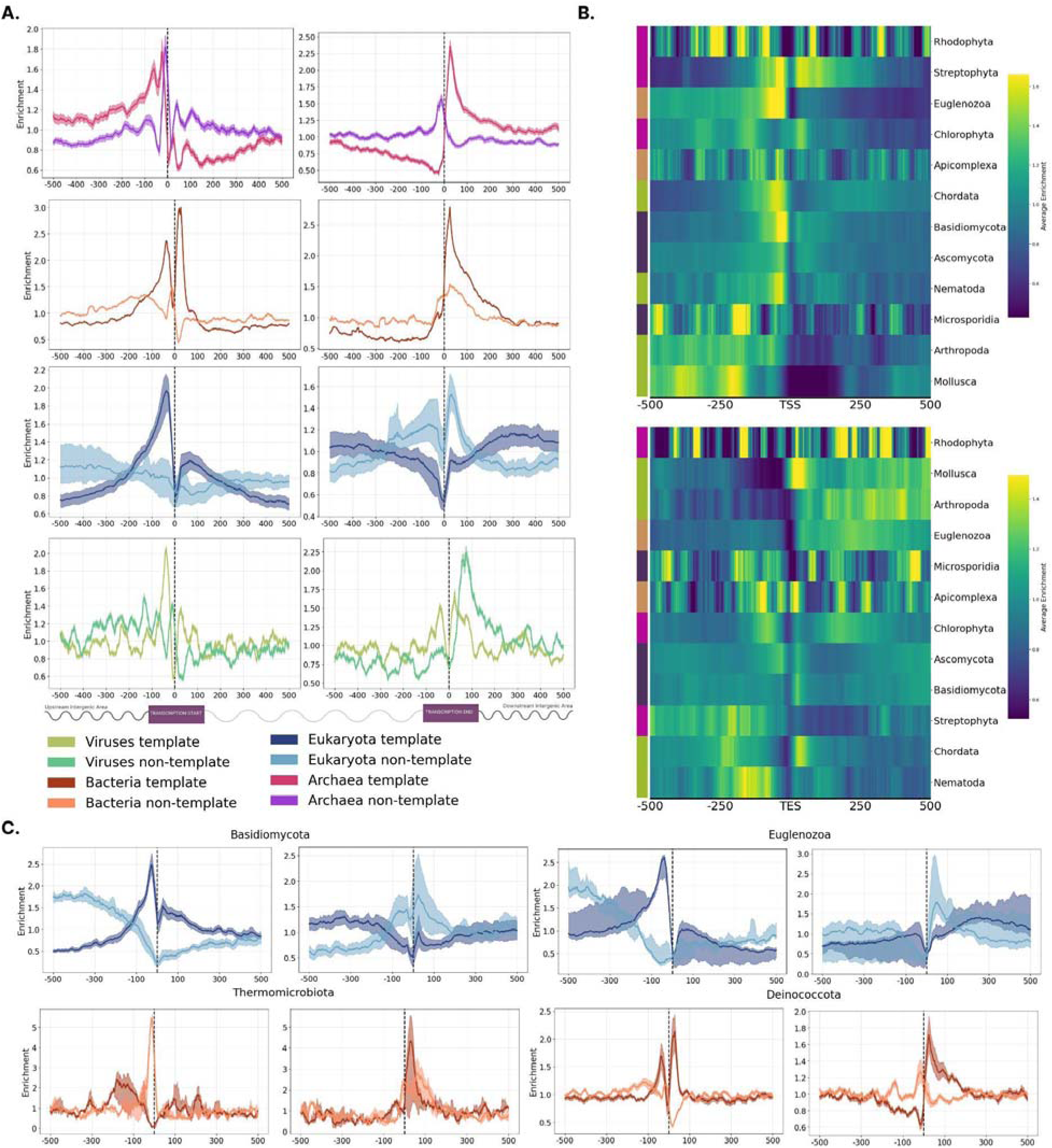
The topography of G4s relative to transcription start and transcription end sites across the tree of life. **A)** G4 distribution across the three domains of life and viruses. Results are shown for the template and non-template strands separately. **B)** Distribution of G4s relative to transcription start and transcription end sites across eukaryotic phyla. **C.** G4 distribution relative to transcription start sites and transcription end sites for two eukaryotic phyla, Basidiomycota and Euglenozoa, and two bacterial phyla, Thermomicrobiota and Deinococcota. Results are shown for the template and non-template strands separately. Error bars represent the 2.5% lowest and 97.5% highest percentile from Monte-Carlo simulations with replacement (N=1,000).

We next investigated how G4 sequences are distributed concerning splice sites in eukaryotic organisms. By comparing the varying frequencies of G4s near splice sites across different taxonomic groups, we sought to understand potential differences in the regulatory roles of G4s in splicing among taxa. We found that G4s display a remarkable enrichment in the phyla of Chordata and Chlorophyta in the intronic area upstream of the acceptor sites and in the intronic area downstream of the donor sites, indicating that G4s regulate the RNA splicing for these eukaryotic phyla (**Figure 5a-b)**. For both G4 detection methods, we observe that there is a roughly 2.5-fold enrichment in the intron of the upstream splice site (3’ss). In the intronic area of the downstream splice site (5’ss), we observe a 3.5-fold enrichment for the regular expression-based consensus motifs and a 2.5-fold enrichment for the G4Hunter-based method. Furthermore, in the aforementioned phyla, significant differences were observed in the corresponding template and non-template distributions (**Figure 5a-b)**. For instance, G4 sequences found in Chordata were predominantly located in the intron preceding the ‘3ss on the template strand, while G4s on the non-template strand were highly enriched in the intronic area downstream of the 3’ss. This indicates that G4s have potential regulatory roles in splicing in Chordata and Chlorophyta, as previously shown from work in humans (14).

**Figure 5:**
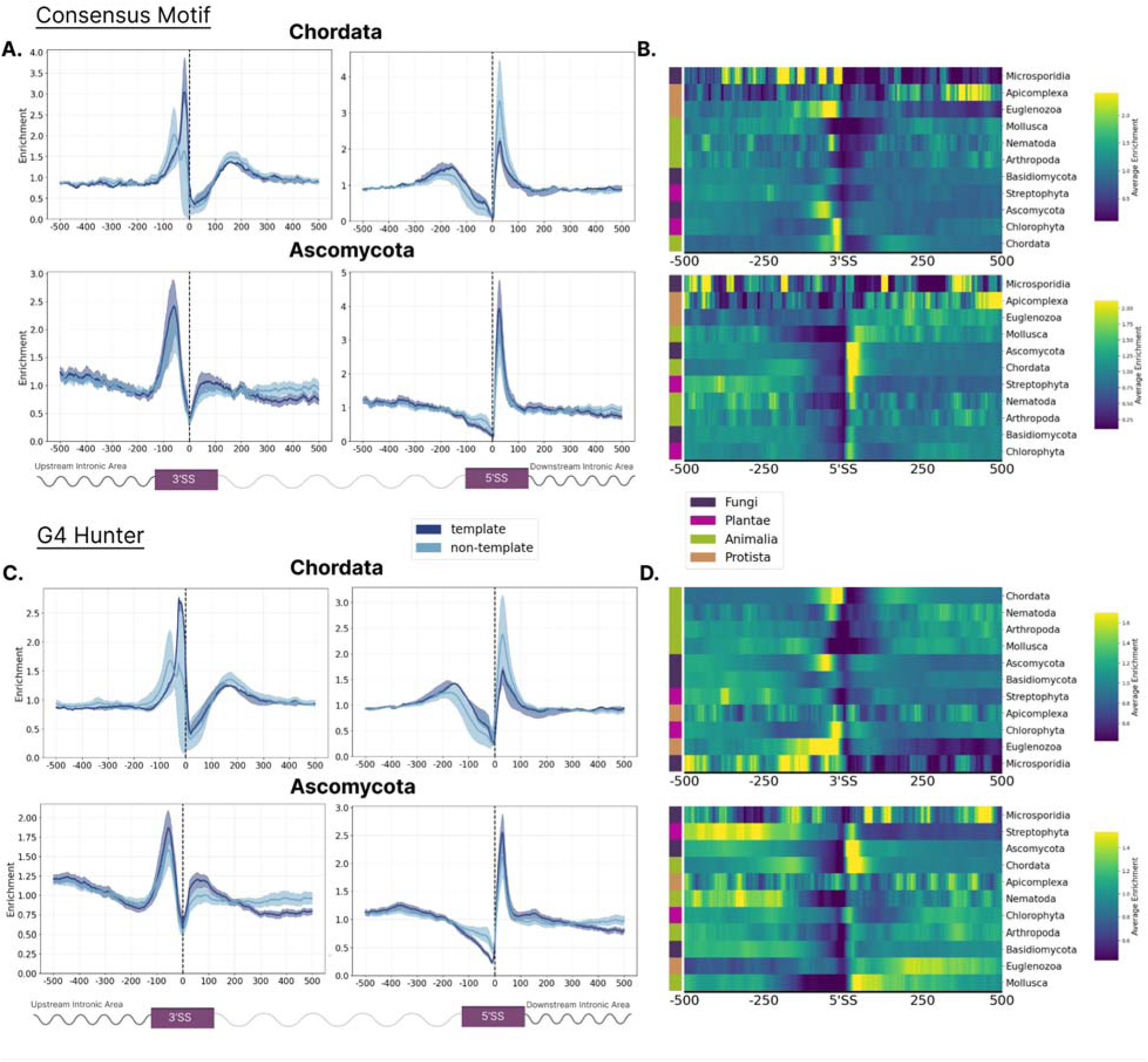
The topography of G4s relative to splice sites in eukaryotic phyla. **A)** Distribution of G4s relative to 3’ss and 5’ss using the consensus G4 motif and the G4Hunter-based algorithm. **B)** Heatmap showing the enrichment of G4s relative to splice sites across different phyla.

### Differences in the intervening loop length in G4 DNA between taxonomies

We also examined the biophysical properties of G4s, including their total length and the length of the intervening loops. The median length of G4s extracted using the regular expression algorithm for the consensus sequence was 25 bps whereas the length distribution of G4s extracted using the G4Hunter tool was 28 bps (**Supplementary Figure 9a**). A G-run constitutes any subsequence of G4 being composed of at least three consecutive Gs, whereas loops are the intervening sequences in a given G4 sequence (**Figure 1a**). We investigated the loop length distribution of all the extracted G4s. Using the G4Hunter algorithm, the most frequent loop length for G4s was nine base pairs, appearing with a frequency of 12%, which also showed a bias towards longer loop lengths. This is opposed to the distribution seen in G4s obtained from the consensus motif, which displayed a more uniform distribution, in which the most common loop length was 17 base pairs, occurring at a slightly higher frequency of 12.3% (**Supplementary Figure 9a**).

As a subsequent step to our analysis, we investigated how the G4s and their associated loop lengths vary across the three domains of life, including viruses. In the consensus motif sequences, we observe that the total G4 length and loop length distributions of G4s originating from eukaryotic or viral organismal genomes show high variance in contrast to archaea or bacteria that are more centralized (**Supplementary Figure 9b)**. Moreover, viruses display a preference for large G4 sequences. In G4s extracted with G4Hunter, we observe that these differences are less pronounced, and the distribution of individual taxonomies is more akin to the distribution containing all organismal genomes. To statistically validate our findings we compared the distributions across the four taxonomies by performing two sample Kolmogorov-Smirnov tests, adjusting for multiple testing. All the distributions were significantly different across both methodologies (p-values<0.0001) (**Supplementary Figure 10**), indicating that the loop length of G4s varies significantly between taxonomies.

### Patterns of G4s in origins of replication and replication strand polarity

We investigated if G4s are differentially found in the forward and reverse strands in bacteria, relative to the origin of replication. To achieve this we generated N=1,000 equal-sized bins per genome, representing the distance from the origin of replication (see Methods). We used only bacteria with a single replication origin, which constitute the vast majority of cases to perform this analysis. In these, there are two diverging replication forks proceeding from the same origin of replication (*oriC*) in opposite directions until they reach the replication termination (*ter*). We mapped G4s in the forward and reverse strands in each genomic bin and observed highly biased distributions when comparing the relative enrichment between the two strands across the genome (**Figure 6a-c; Supplementary Figure 11**). These biases reflect a preference for G4s in the leading strand and a dearth in the lagging strand. At individual phyla, we observe strong biases in G4 preference in Baciliota and Pseudomonadota, but not in Cyanobacteria (**Figure 6a-c; Supplementary Figure 11**).

**Figure 6:**
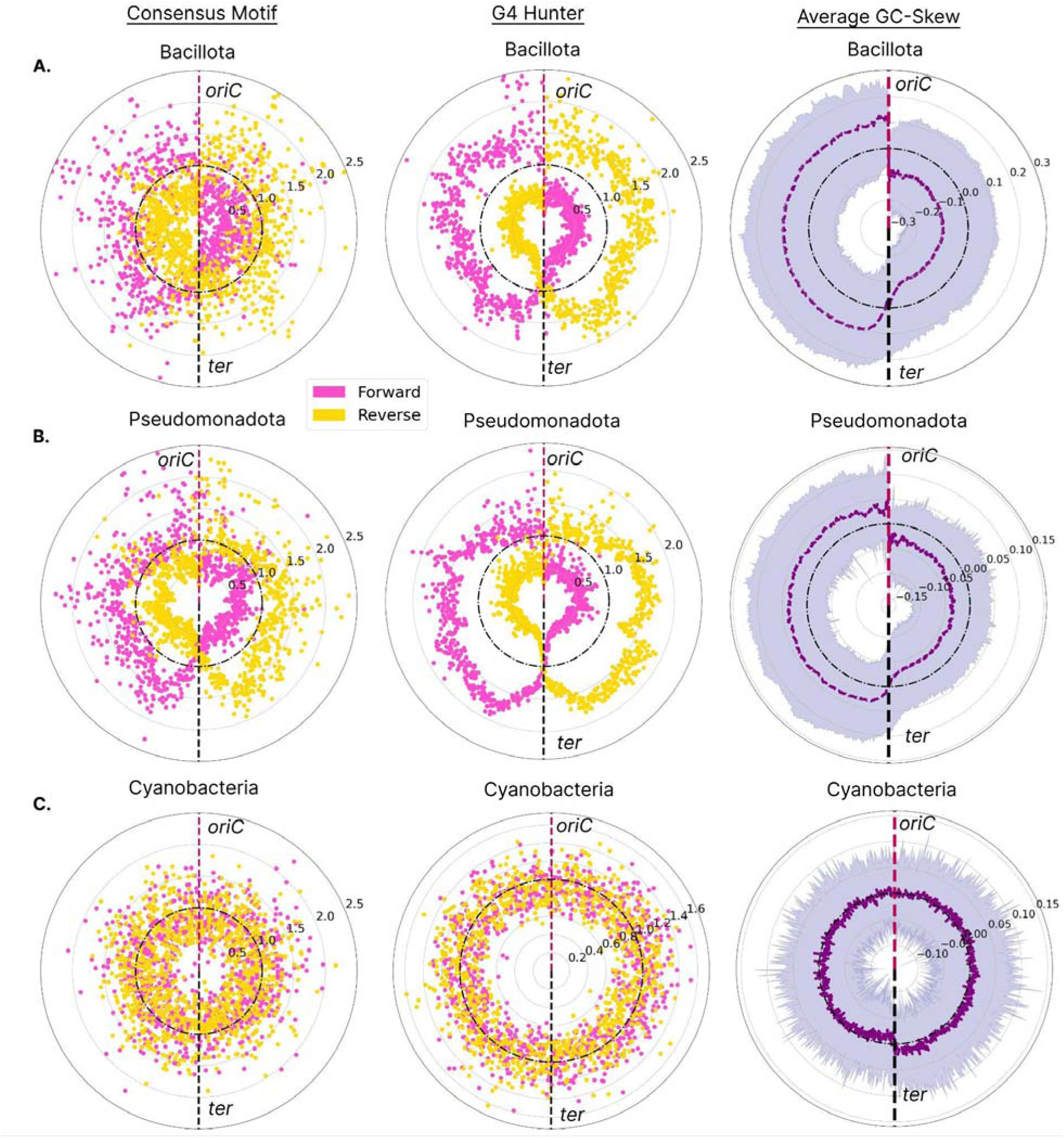
G4 distribution patterns relative to replication origin in bacterial phyla. Results shown for **A)** Bacillota, **B)** Pseudomonadota, **C)** Cyanobacteria. Enrichment of G4s in forward and reverse strand orientation are shown in yellow and pink, respectively. Results are shown for G4 Hunter-based and Consensus Motif-based algorithms. We discretized the circular bacterial chromosomes in 1001 bins, relative to oriC, for each calculating the total number of G4 sequences divided by the total number of G4s that span the whole chromosome, to estimate the local enrichment of G4s. The average GC skew is also calculated and shown in purple.

To further evaluate these differences, we calculated the average GC skew levels (GC skew = (G − C)/(G + C)) in the different parts of the genomes of each organism in these phyla (80). Guanines are more abundant in the leading strand, and therefore negative GC skew scores are linked to the leading strand (80, 81). Our observations indicate that phyla exhibiting pronounced GC skew biases between leading and lagging strands tend to have differences in the levels of G4s between the leading and lagging orientations. Conversely, in phyla such as Cyanobacteria, where the GC skew is less pronounced, there is less disparity in the G4 frequency between the strands. Consequently, the distribution of G4s between leading and lagging strands in these phyla tends to be more uniform. We conclude that the distribution of G4s across bacterial genomes of specific phyla can be profoundly biased towards the leading strand, dictating their topology.

### Experimental validation of predicted G4s

We selected a subset of potential G4 DNA-forming sequences to experimentally validate their ability to adopt G4 DNA *in vitro* using circular dichroism spectroscopy, UV-melting, and fluorescence measurements. These included G4 DNA-forming sequences that were highly prevalent across species (found in at least 2,000 species), and which were either only detected by G4-hunter or the consensus regex-based G4 algorithm (**Supplementary Table 2)**. We conducted several spectroscopic analyses on each of the seven candidates to validate the formation of G4 structures. Initially, we utilized circular dichroism spectroscopy and UV-melting assays on DNA oligonucleotides that contained the potential G4 DNA-forming sequences. These tests were done in the presence of either lithium ions (Li^+^), which do not stabilize G4s, or potassium ions (K^+^), which do, to assess the potential and stability of G4 formation. Overall, a higher signal was found under K^+^ conditions in all CD spectra, suggesting the presence of G4 structures in all sequences tested (**Figure 7a**). The CD spectra also reveal different G4 topologies formed in the different sequences, including parallel, antiparallel, and hybrid topologies (**Figure 7a**). In support of the presence of G4s, our findings suggest that all candidates formed thermostable G4 structures under K^+^ conditions, which was confirmed by the hypochromic shift observed in all UV spectra, with the melting temperatures determined to be within a range of 40 to 80 °C (**Figure 7b**). Additionally, we employed fluorescence-based techniques, including the use of N-methyl mesoporphyrin IX (NMM) ligand-enhanced fluorescence and ISCH-oa1-enhanced fluorescence experiments (**Figure 7c–d**). In all cases, a higher fluorescent intensity was observed under K^+^ conditions compared to Li^+^ conditions, which indicates the formation of G4 structures in all selected sequences.

**Figure 7:**
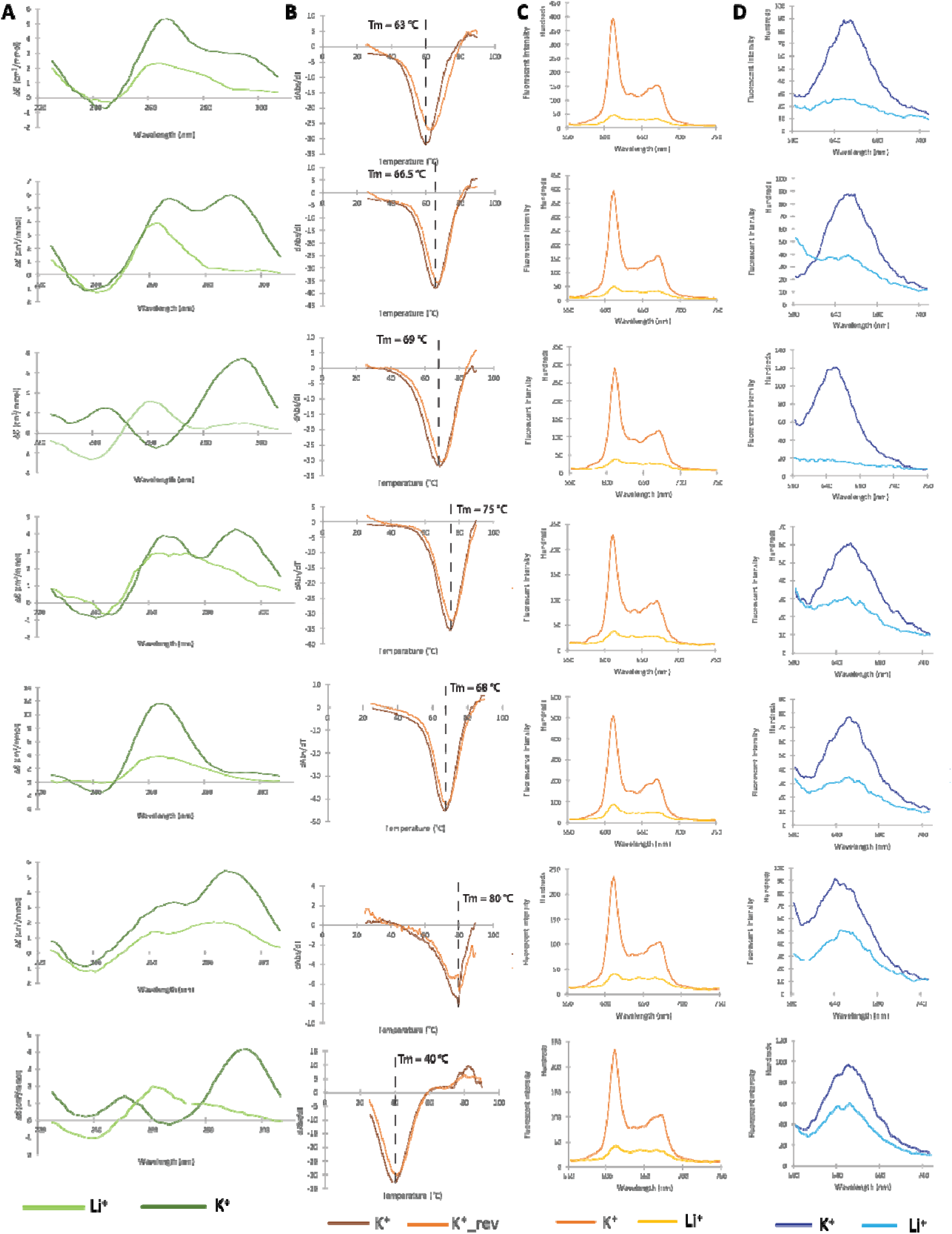
Experimentally validated G-quadruplex candidates. G4 ligand-induced G4 spectroscopic analysis reveals the G4 DNA structure formation in selected sequences from the dG4 database. **A)** CD spectra in the presence of K^+^ and Li^+^. The overall higher signals under K^+^ conditions verify the presence of dG4 in all sequences. The positive and negative peaks at different wavelengths suggest different topologies were formed in different sequences, including parallel (negative peak at 240 nm, positive peak at 260 nm), anti-parallel (negative peak at 240 nm, positive peaks at 240 nm and 295 nm), and hybrid (negative peak at 240 nm, positive peaks at 260 nm and 295 nm) topologies. **B)** UV melting spectra in the presence of K^+^. The hypochromic shift observed at a wavelength of 295 nm is consistent with the presence of dG4 structures, with the melting temperature determined as the maximum negative value of the hypochromic shift. **C)** NMM, and **D)** ISCH-oa1 enhanced fluorescence spectroscopy in the presence of K^+^ and Li^+^. The higher fluorescent signals under K^+^ conditions illustrate the dG4 structure formation in all sequences.

### Derivation of G4 clusters

We next performed a clustering analysis across all the identified G4s in all genomes and grouped G4 sequences based on sequence similarity. After accounting for overlapping G4 loci, the final set used for clustering comprised 118,445,719 G4s, of which 88,139,047 came from the G4hunter-based algorithm, 8,571,114 were derived from the regex-based algorithm, and 21,735,558 were unique representatives of sequences reported by both methods. A total of 6,819,259 G4 sequence groups of one or more members were produced; 319,784 of which had 20 or more G4 sequences and were selected as the final cluster dataset.

We observe that the majority of G4 clusters are of eukaryotic origin, followed by bacterial origin (**Figure 8**). When we further separate the G4 clusters by eukaryotic taxonomic subdivisions, we find that non-mammalian vertebrates and plants have the largest number of G4 clusters. Notably, the vast majority of G4 clusters are domain-specific, i.e. come from sequences found in a specific domain of life (e.g. bacteria), and only a significantly small we portion represents G4s found in multiple domains (e.g. bacteria alongside archaea), with only 135 G4 clusters containing sequences from all four organism groups (bacteria, archaea, eukaryota and viruses). The distribution of sequences based on their detection method shows that the majority of the clusters (263,885) contain exclusively sequence from G4hunter (G4hunter-based), followed by 65,868 clusters with sequences from both methods (mixed clusters) and, finally, 250 clusters exclusively containing regex-derived results (regex-based). The number of cluster members varies substantially, with most frequent having 20-50 G4 sequence members for the G4-Hunter-based,regex-based, and mixed clusters.

**Figure 8:**
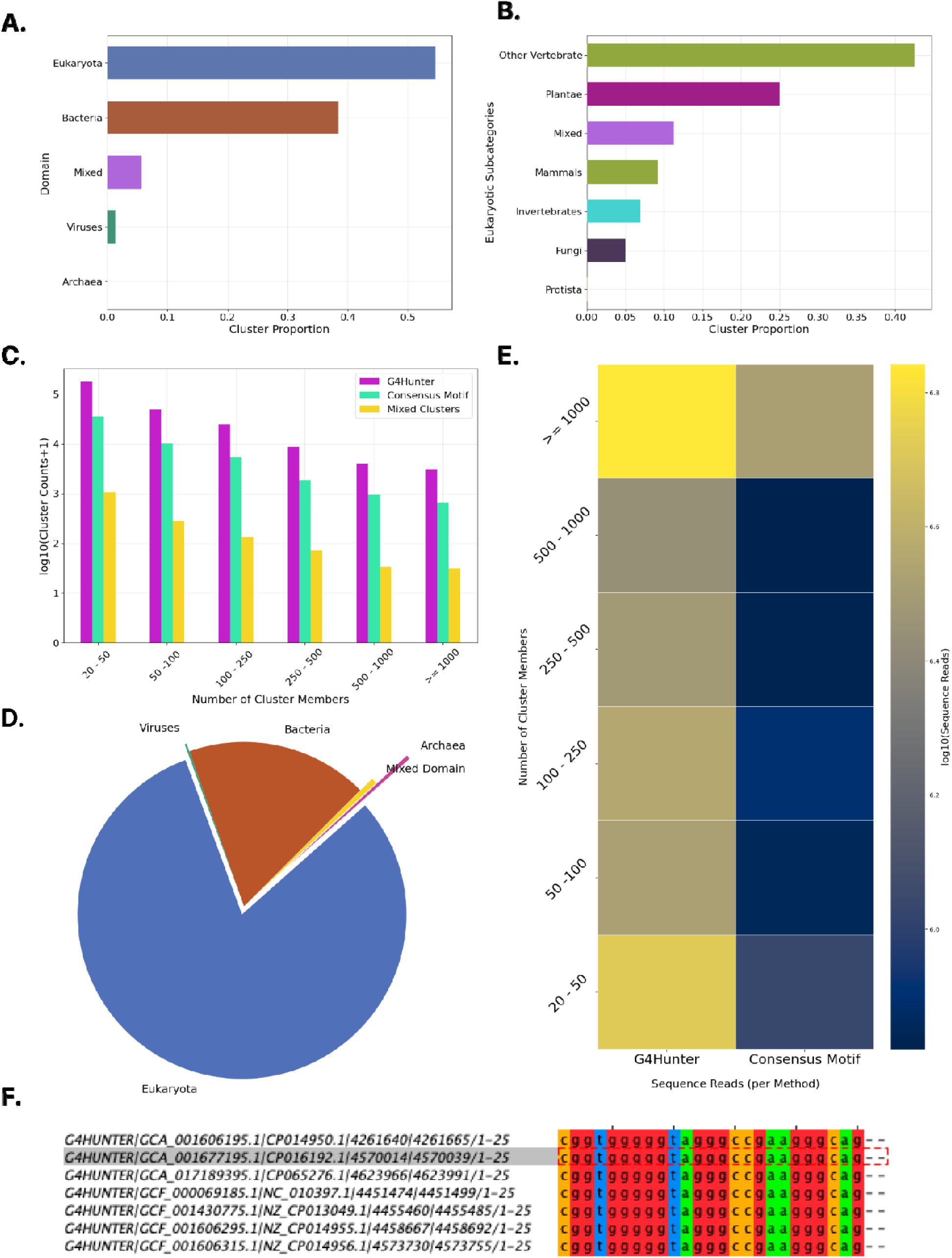
Clustering analysis of G4 sequences. **A)** Proportion of total G4 clusters identified in each of the three domains of life and viruses, as well as those clusters that were found in at least two of these (mixed). **B)** Proportion of clusters observed in each of the kingdoms of life. **C)** Number of clusters based on the number of G4 sequences being members. Results are shown for the consensus motif algorithm and the G4 Hunter algorithm. **D)** Origin of G4 clusters as a proportion of total sequences. **E)** Number of cluster members. Sequence reads were defined as the total number of G4s detected within the indicated cluster. **F)** Example of G4 cluster showing the loci of the G4 sequences, and the associated sequence alignment.

### The Quadrupia website and web-interface

The Quadrupia website contains interactive tables and drop-down menus that enable the selection of potential G-quadruplex DNA-forming sequences based on species names or accession IDs (**Figure 9a**). The data contained in Quadrupia can be accessed through the Browse menu located on the Quadrupia navigation bar. A user can navigate the database for different genomes as well as taxonomic groups of bacteria, eukaryotes, archaea, and viruses. In the database, each genome is represented using its NCBI genome accession as its primary identifier (e.g. GCA_030914265.1) The user can also specify between genomes or use a combination of both criteria. The Quadrupia browse page displays a list of genomes that can be parsed to match the selected filters (**Figure 9b**). The user can further refine the search by opting to select specific species using the NCBI accession, or the species name, which takes the user to the specific genome entry page (**Figure 9c**). On this page, the user can view the G4s, from each of the two detection methods, associated with the selected genome. The information on this page includes the species name, the taxonomic ID, the taxonomic group, the assembly accession with an embedded link to the NCBI webpage, the assembly name, the sequencing level, and the source database (**Figure 9c**). On the same page, the total number of G4s identified with each of the two methods is provided as well as the top 50 longest G4s for each of the two methods.

**Figure 9:**
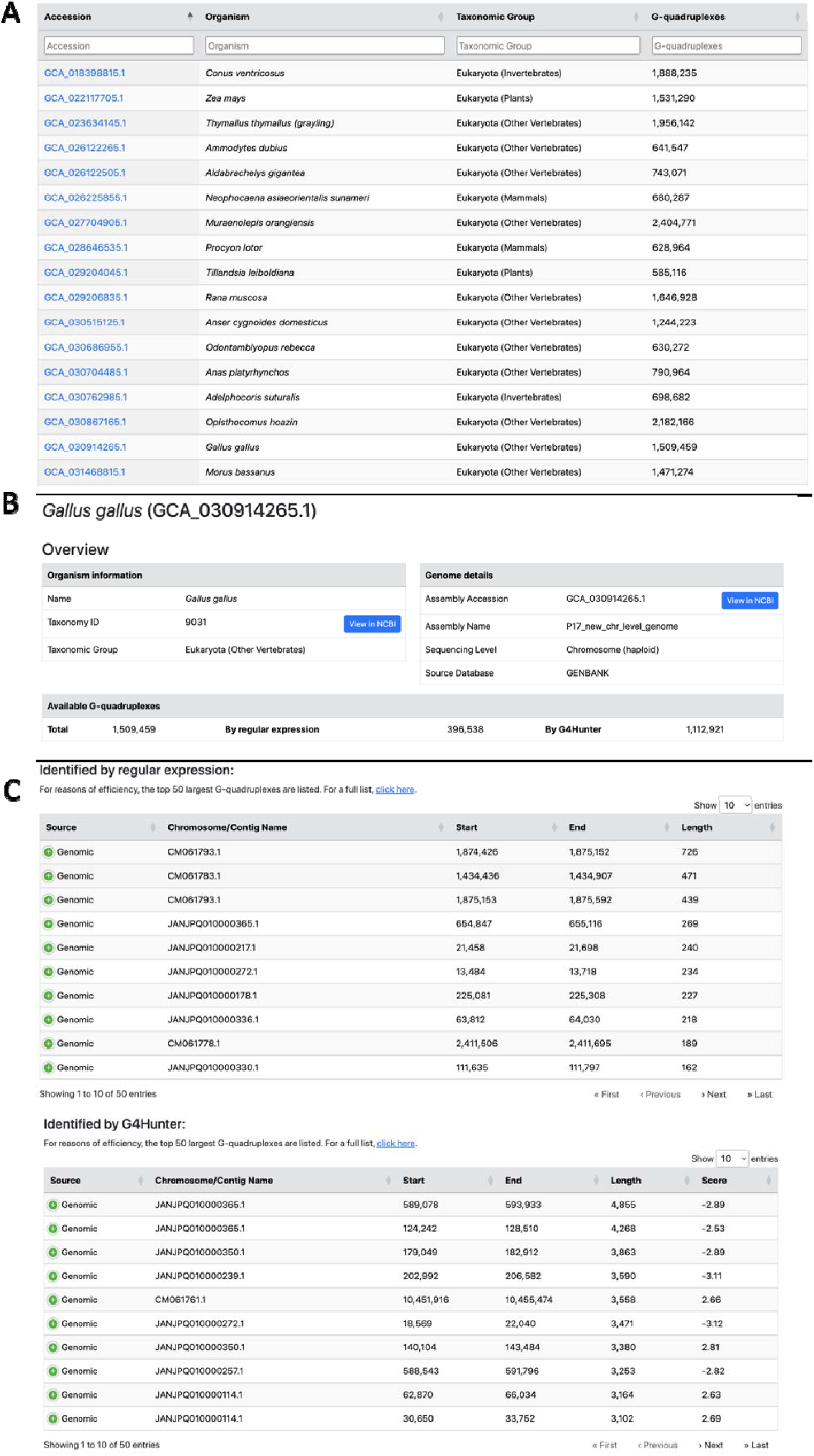
Quadrupia browse pages for reference genomes. **A)** A table of species entries. By default 20 entries/page are shown with the following columns: Accession, Organism, Taxonomic Group, and number of G4s. **B)** The Quadrupia Genome Entry page. An example is shown for the genome of *Gallus gallus* (GCA_030914265.1). The page includes the name of the organism, the taxonomy ID, the taxonomic group, the assembly accession, the assembly name, the sequencing level, and the source database. **C)** G4s identified using regular expressions and G4Hunter. The output is sorted by the length of the G4 and the table displays the source, chromosome/contig name, the start and end coordinates of the G4, its length, and for G4Hunter, the G4Hunter score. The table is sortable and the results can be downloaded.

Similar to the genomes, the database allows navigating the collection of G4 clusters (**Figure 10**). Each cluster in the database is represented by a unique identifier in the form of G4-XXXXXXX (e.g. G4C-0084453). The clusters are accessible through a separate browser, available from the top navigation menu. Each cluster can be viewed by its distinct Cluster Entry page, and is accompanied by a number of annotation data, including taxonomic distribution, a multiple sequence alignment (MSA) and its derived Hidden Markov Model (HMM), and a centroid, representative sequence. The MSA and HMM are available for visualization in the Cluster Entry page through an interactive alignment browser and a sequence logo viewer, respectively. In addition, the MSA and HMM raw files are available for download through buttons at the top of the page. Finally, in cases of clusters that have sequence homology to experimentally determined G4 structures, external links to the latter are given for the PDB (71) and OnQuadro (61) databases, and the top homolog structure is available for visualization through an interactive 3D structure viewer.

**Figure 10.**
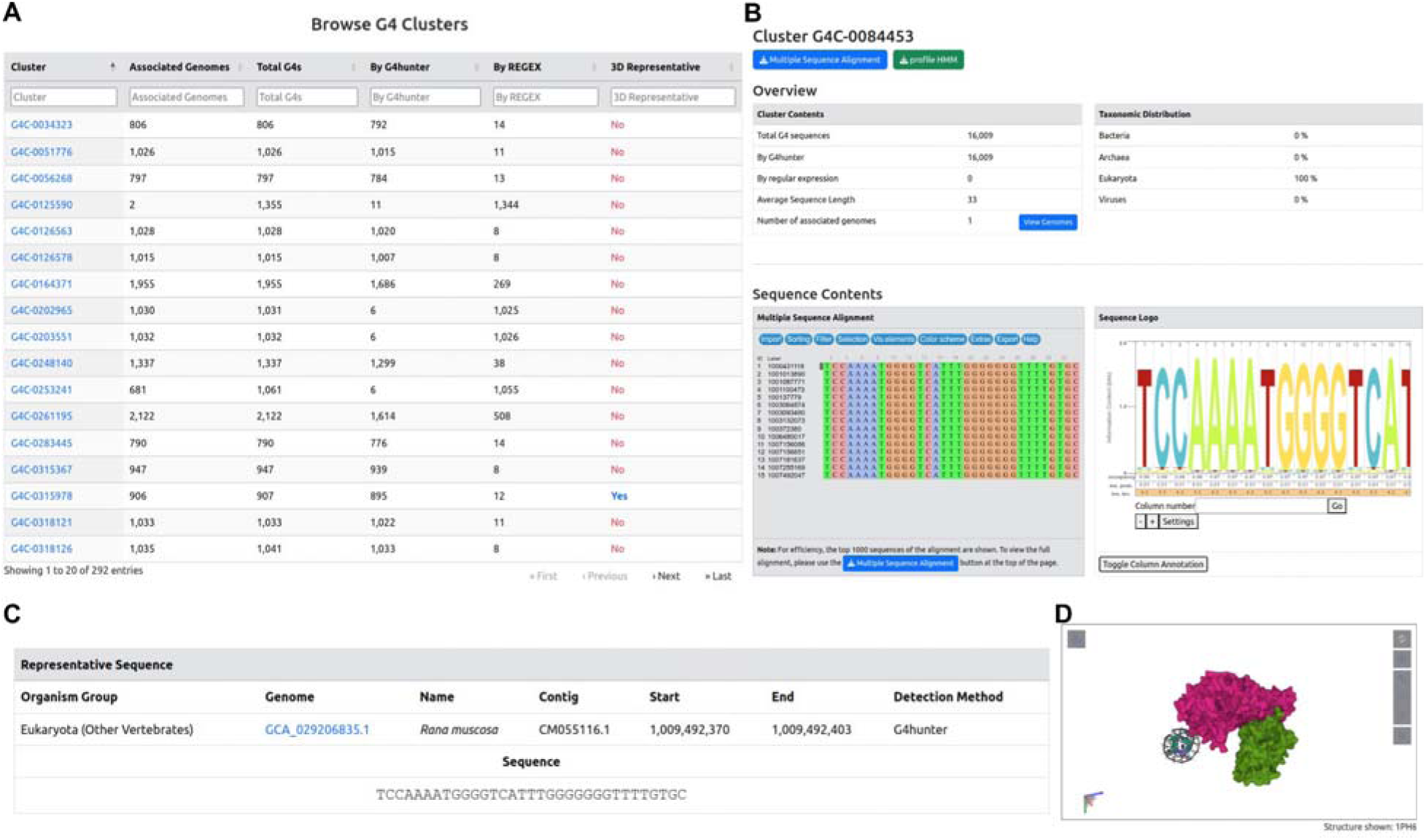
G4 cluster search and visualization. **A)** The G4 cluster browser. Each cluster is identified by a unique identifier and is annotated by the number of associated genomes, the number of G4 sequences (total, by regular expression or G4hunter), and, finally, the presence or absence of a 3D representative. **B-D)** An example G4 cluster entry page (ID: G4C-0084453). The entry contains the basic cluster metadata, as well as the taxonomic distribution of its contents **(B)**. In addition, interactive viewers are given for the cluster’s MSA, HMM (in the form of a sequence logo), representative sequence **(C),** and, if available, 3D structure representative conformation **(D)**.

The database is also searchable with a Sequence Search Option or an Advanced Search Option. The Advanced Search page allows performing complex queries against the database by combining multiple fields, such as the taxonomy, number of G4 sequences and sequence detection method. The Sequence Search Option (**Figure 11**), in turn, allows users to perform the following queries: i) search DNA/RNA sequences against G4 sequences, performed using pairwise alignments with BLAST (blastn), ii) search sequences against the collection of G4-clusters, using HMM-based queries with HMMER (nhmmer) and, iii) search for hits to custom sequence motifs, either in the form of regular expressions or as IUPAC sequences Finally, Quadrupia provides a Downloads page, which enables users to download the complete datasets and incorporate them in their analyses.

**Figure 11:**
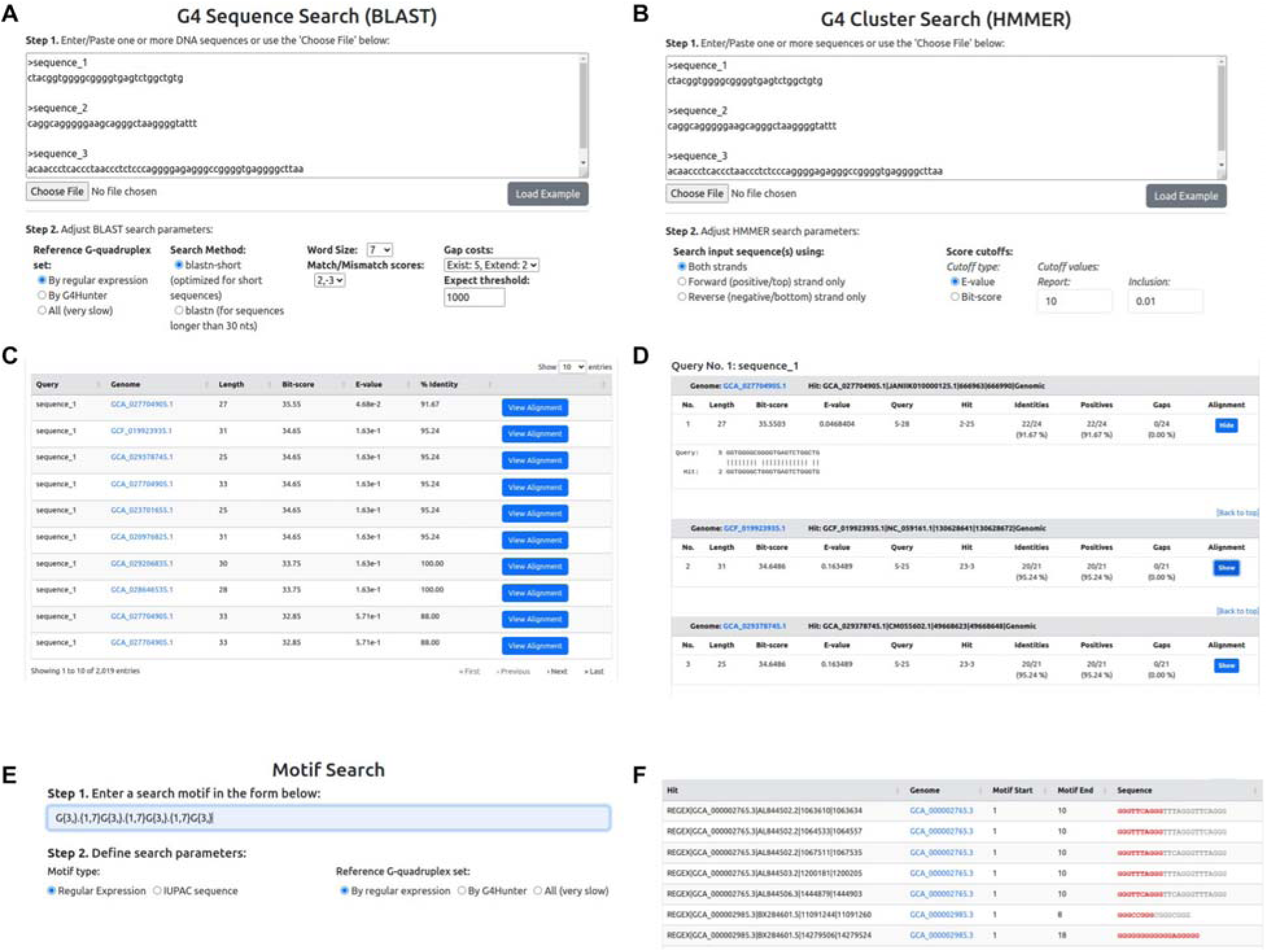
The Sequence Search Options. A) G4 Sequence Search input page. Search options include defining the search algorithm (blastn or blastn-short), setting the word size and defining the match/mismatch scores and gap penalties. B) G4 Cluster Search. Users can choose which DNA strand to include in the search and define cutoffs for the E-value and Bit-score metrics. C-D) Example BLAST results. An overview (C) of the results presents the basic search statistics (alignment length, Bit-score, E-value and % sequence identity), while the detailed results (D) give the full BLAST output, including a visualization of the aligned sequences. E-F). Input form (E) and example results (F) of the Motif Search functionality. Sequence motifs can be given as regular expression patterns or IUPAC sequence stretches. The results include all G4 sequences that match the submitted motif.

### Documentation and Help pages

The Quadrupia website has a “Documentation” page which provides background information on G4s and a “Help” page about how to navigate through the G4 data, the different species as well as how to download the reference maps.

## Discussion

We have performed a systematic characterization of potential G4 DNA-forming sequences across organismal genomes spanning the tree of life, including the three domains of life and viruses, diverse kingdoms, and phyla. We introduce Quadrupia, a comprehensive database dedicated to the study and analysis of G4 DNA. Our database encompasses genome-wide maps of G4 sites for 108,534 organisms, far exceeding the number of species covered in comparison to any other G4-associated database. The database incorporates searchable options using motif-based or BLAST-based searchers. We also perform large-scale analyses on the distribution and topology of G4s across these organismal genomes. We find that eukaryotes are the domain with the highest G4 density; nevertheless, there are large differences at the phylum level, with the phyla with the highest G4 density belonging to different domains of life and kingdoms, suggesting that G4s are genomic elements with high turnover between taxonomic subgroups.

The prevalence of G4 DNA-forming sequences in organisms across the tree of life suggests that G4s are functional elements across organismal genomes. However, in our analysis, G4s were unevenly distributed across the three domains of life and viruses. Their positioning in genomic loci associated with regulatory roles, such as in the broader promoter regions, near transcription end sites, and in intronic regions close to splice sites, highlights their involvement in mediating a spectrum of important biological functions and mechanisms. We also observe that these enrichments in regulatory regions can be taxonomy-specific. It is possible that due to distinct environmental influences, and different organism needs, G4s have dynamically evolved.

The diversity in which G4s are distributed across different organisms, kingdoms, and phyla is suggestive of their intrinsic regulatory roles for various molecular functions and biological mechanisms. Notably, the phylum of Deinococcota displayed the highest G4 density using the two G4 detection algorithms. Moreover, significant enrichment of G4s was detected upstream of the TSS and downstream of the TES in the Deinococcota phylum. This is in accordance with a previous work (82), in which the bacteria belonging to the Deinococcota phylum are highly resistant to environmental hazards, with high survival rates when exposed to gamma or UV radiation, and in which G4s have regulatory and radioresistant functions (82). The phylum of Peploviricota displayed a high density of G4s, appearing second in consensus G4 motifs and fourth using the G4Hunter algorithm. Viruses belonging to this phyla also constituted the majority of the top 100 viruses with the highest G4 densities. This viral phylum consists solely of a single order, Herpesvirales, an order of double-stranded DNA viruses, which encompasses a variety of viral species that affect animal hosts. Previous work shows that the viral cycle of the herpes simplex virus-1 (HSV-1) is regulated by G4s. Due to the central role of G4s in the viral cycle, G4s have been used as drug targets for antiviral activity (83). Additionally, we observe that viruses with eukaryotic hosts have significantly higher G4 densities compared to viruses with prokaryotic hosts. This might suggest that eukaryotic viruses evolved to acquire higher G4 densities to adapt to the G4-rich environment in their eukaryotic hosts. However, due to the large variation in G4 densities at the phyla level, within the same domains, further studies need to be done between viruses and their hosts at lower taxonomic levels.

Other significant G4 densities are found in various plant phyla such as Chlorophyta, which display a high overall G4 density and a significant enrichment in regulatory regions such as TSSs, TESs and splice sites. These positional tendencies of G4s in central regulatory regions are not limited to bacteria, viruses, or plants. We highlight a significant density of G4s in various eukaryotic kingdoms such as fungi and protozoa (**Figure 2**). Several Fungi phyla such as Microsporidia, Basidiomycota, and Ascomycota display a high density of G4s in their genome and an enrichment of G4s within the broader promoter regions,(**Figure 4**). For *Aspergillus fumigatus*, of the phylum Ascomycota, G4s have been associated with genes, involved with virulence, and implicated in drug resistance, and could be used for the identification of novel antifungal targets (84). Similar results have been identified amongst Protozoa, with the phylum of Euglonozoa displaying similar G4 enrichment in regulatory areas.

In bacteria, we find that G4s are preferentially found in the leading strand during replication. However, we also observe that this strand asymmetry of G4s is phylum-specific and is associated with the GC skew levels. There is evidence that there are more guanines than cytosines in the leading strand in bacteria (80), which could result in a higher likelihood of formation of G4s. We also observe that certain phyla, such as Cyanobacteria and Deinococcota lack these biases, and do not have strong GC skew biases. Previous research has shown that the GC skew is less pronounced in Cyanobacteria (85, 86), which is consistent with our findings and which could therefore account for the absence of strand asymmetry between leading and lagging strands in the distribution of G4s. G4s can be barriers to replication fork progression (87), and the observed preference of G4s for the leading orientation could influence genome stability. Future work is required to decipher the extent to which the observed differences in the frequency of G4s on the leading and lagging strands impact the mutation rate in bacterial genomes.

G4s are elements with increased mutagenicity and high turnover rate (4, 26, 88, 89). We observe that across taxonomic groups, the G4 density has extreme variation, suggesting that they are plastic genomic elements driving evolution (79, 89). We also find that functional genomic sub-compartments show large deviations in G4 density between taxonomic sub-groups, which likely reflects the emergence of different functionalities when comparing taxonomic clades. These findings are further supported by previous results indicating that the roles of G4s in splicing have emerged during eukaryotic evolution (14) and previous findings from 37 species showing that the density of G4s has increased in higher eukaryotes (90). We also note that even though this trend exists in bacteria and eukaryotes, for archaea this trend was observed only for consensus G4 motifs, whereas viruses did not display this property.

To conclude, Quadrupia stands out as a multifaceted and user-friendly database, offering G4 sequences and their annotations across species and taxonomic classifications as well as several search options. Further work is required to follow individual G4s in evolutionary lineages and to understand the functional roles of conserved and recently evolved G4s. Such work can include comparative genomics and phylogenetic analyses, enabling additional insights into adaptive strategies and evolutionary constraints. Finally, previous work has indicated differences in the stability of G4s between species (90). Future studies are required to examine differences in the formation likelihood, kinetics, and stability of G4s between disparate taxonomic groups and their implications.

## Code Availability

The GitHub code is provided at: https://github.com/Georgakopoulos-Soares-lab/g4_analysis

## Acknowledgements

We would like to thank Kateryna Makova and Kaivan Kamali for helpful comments.

## Supplementary Figures

**Supplementary Figure 1:**
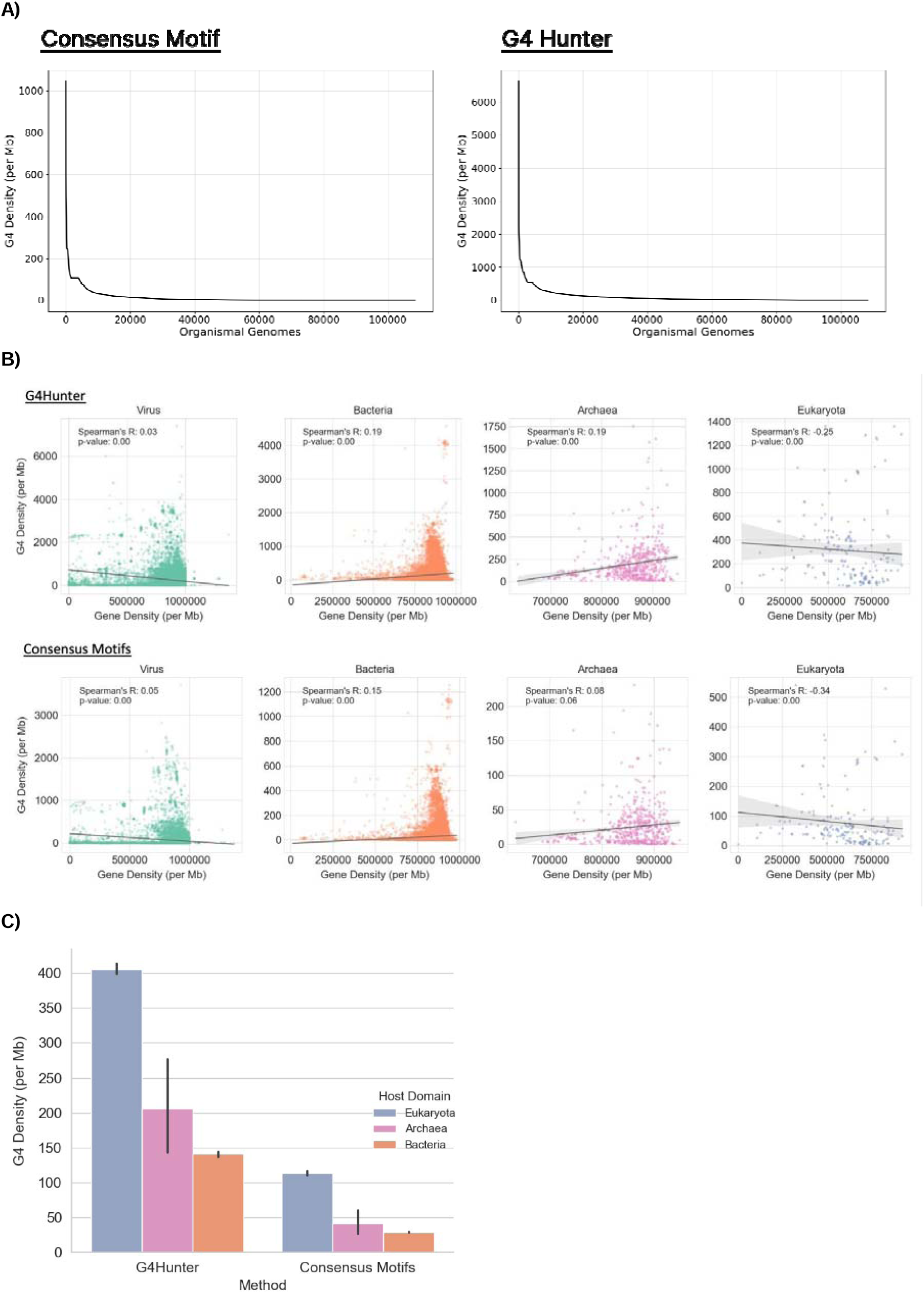
Distribution of potential G4 DNA-forming sequences across organismal genomes and taxa. **A)** G4 density per Mb per organismal genome. **B)** Association between the gene density of a genome and the G4 density, stratified by taxonomic subdivision in the three domains of life and viruses. **C)** Association of G4 densities of viral genomes with their hosts.

**Supplementary Figure 2:**
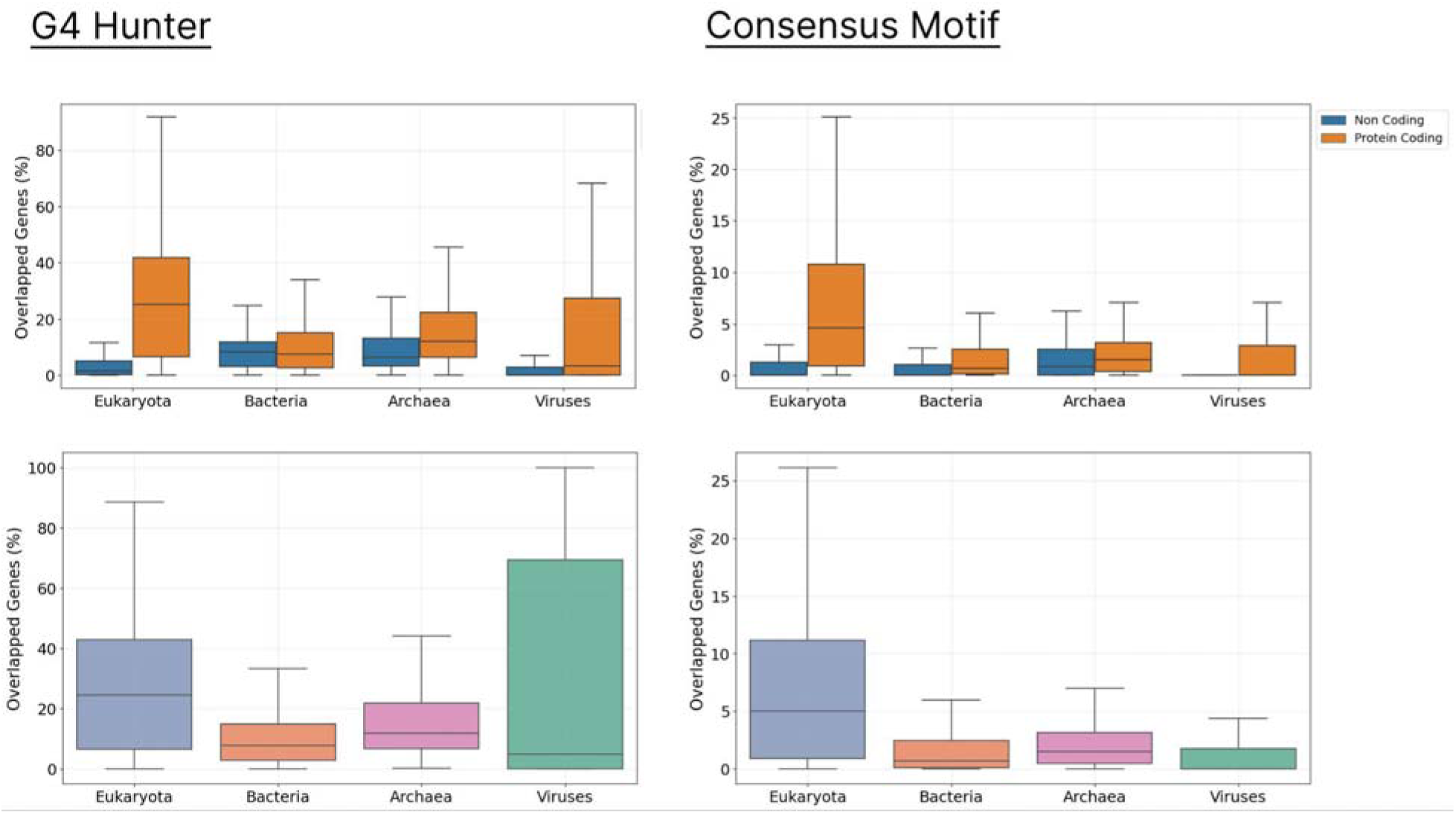
Proportion of G4s overlapping genes.

**Supplementary Figure 3:**
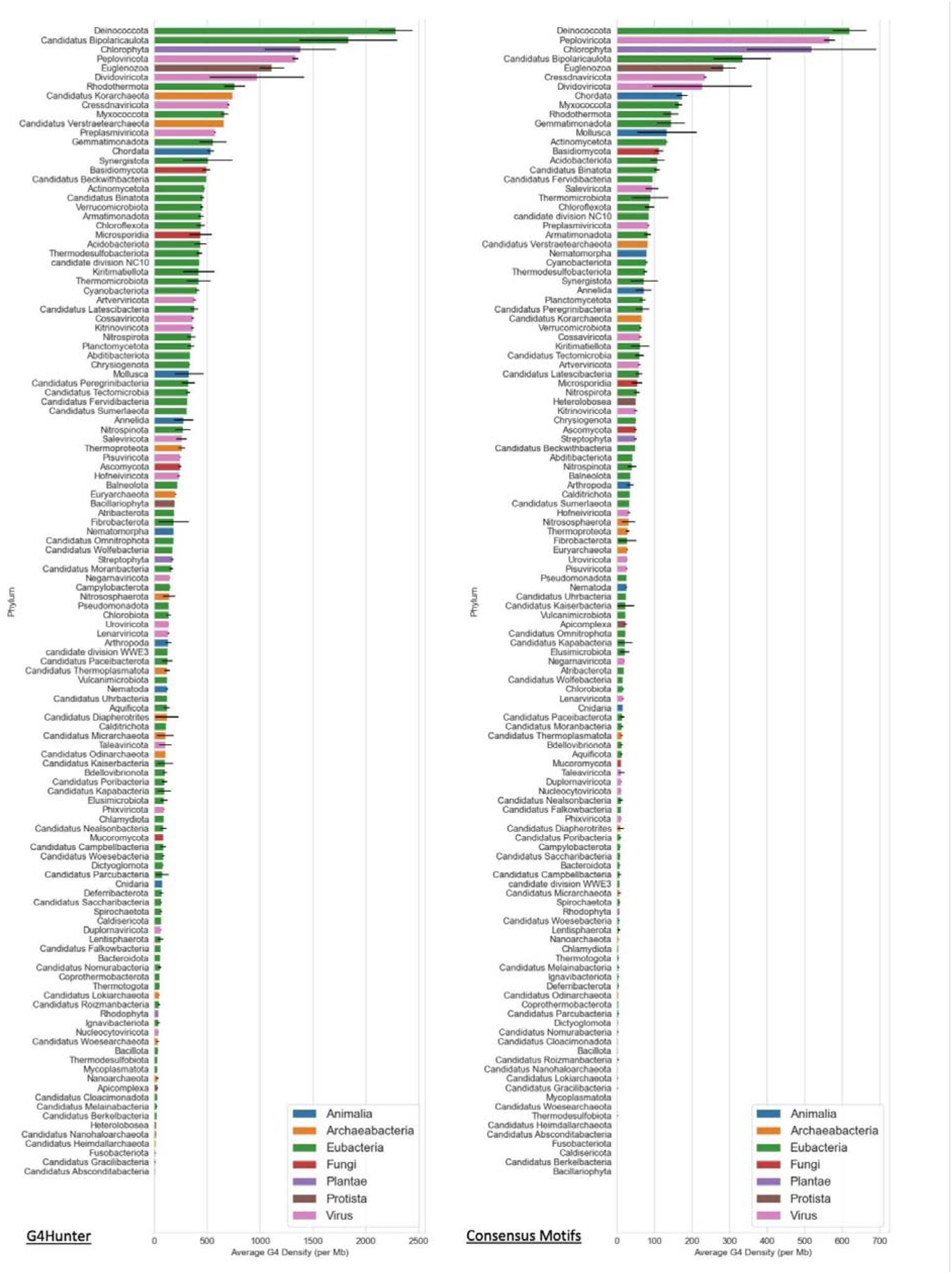
Density of G4s in each phylum. Error bars show standard deviation from the mean. All phyla are displayed.

**Supplementary Figure 4:**
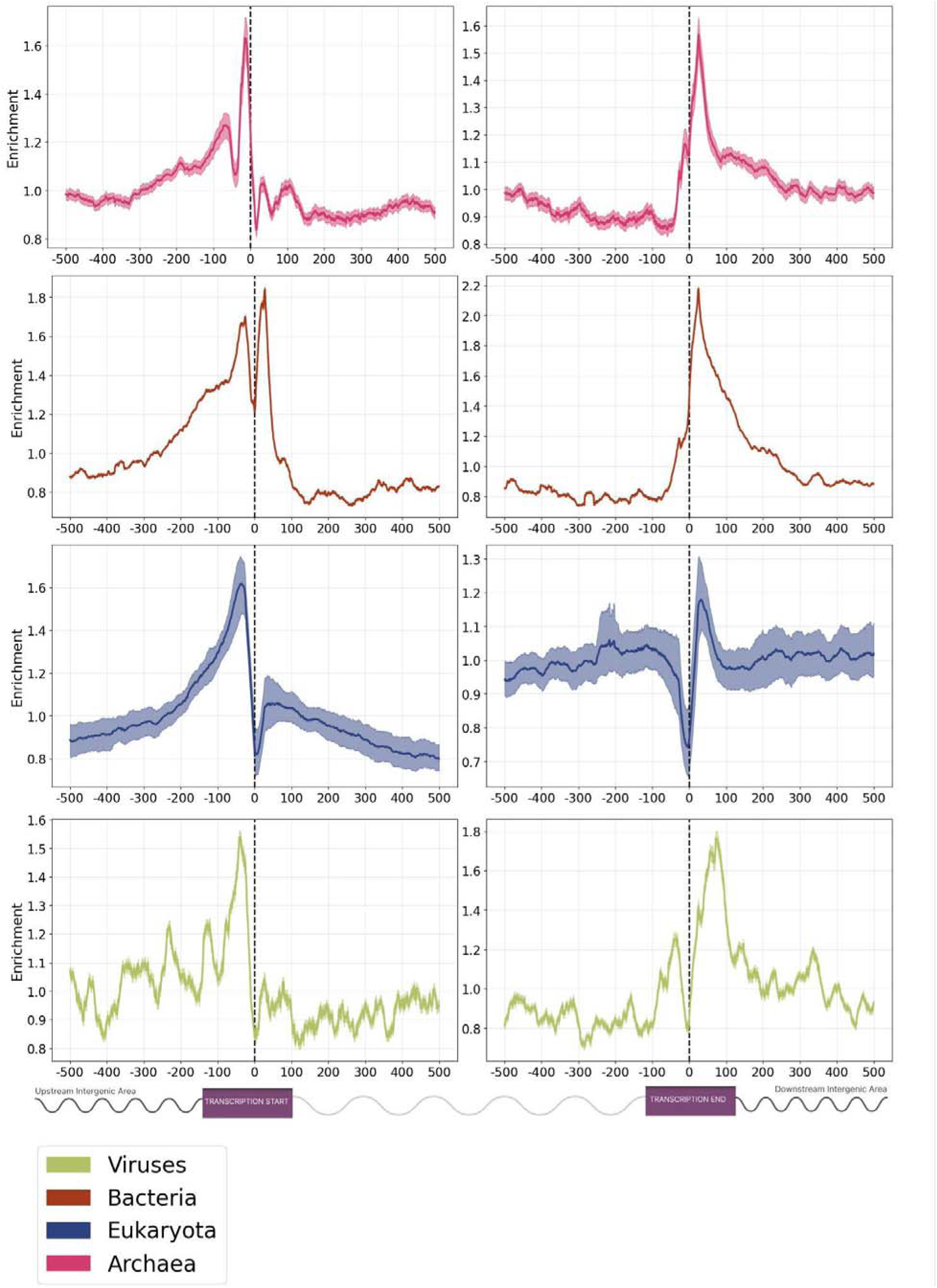
The topography of G4s relative to transcription start and transcription end sites across the tree of life. G4 distribution across the three domains of life and viruses derived using the G4Hunter algorithm. Confidence intervals represent the 2.5% lowest and 97.5% highest percentile from Monte-Carlo simulations with replacement (N=1,000).

**Supplementary Figure 5:**
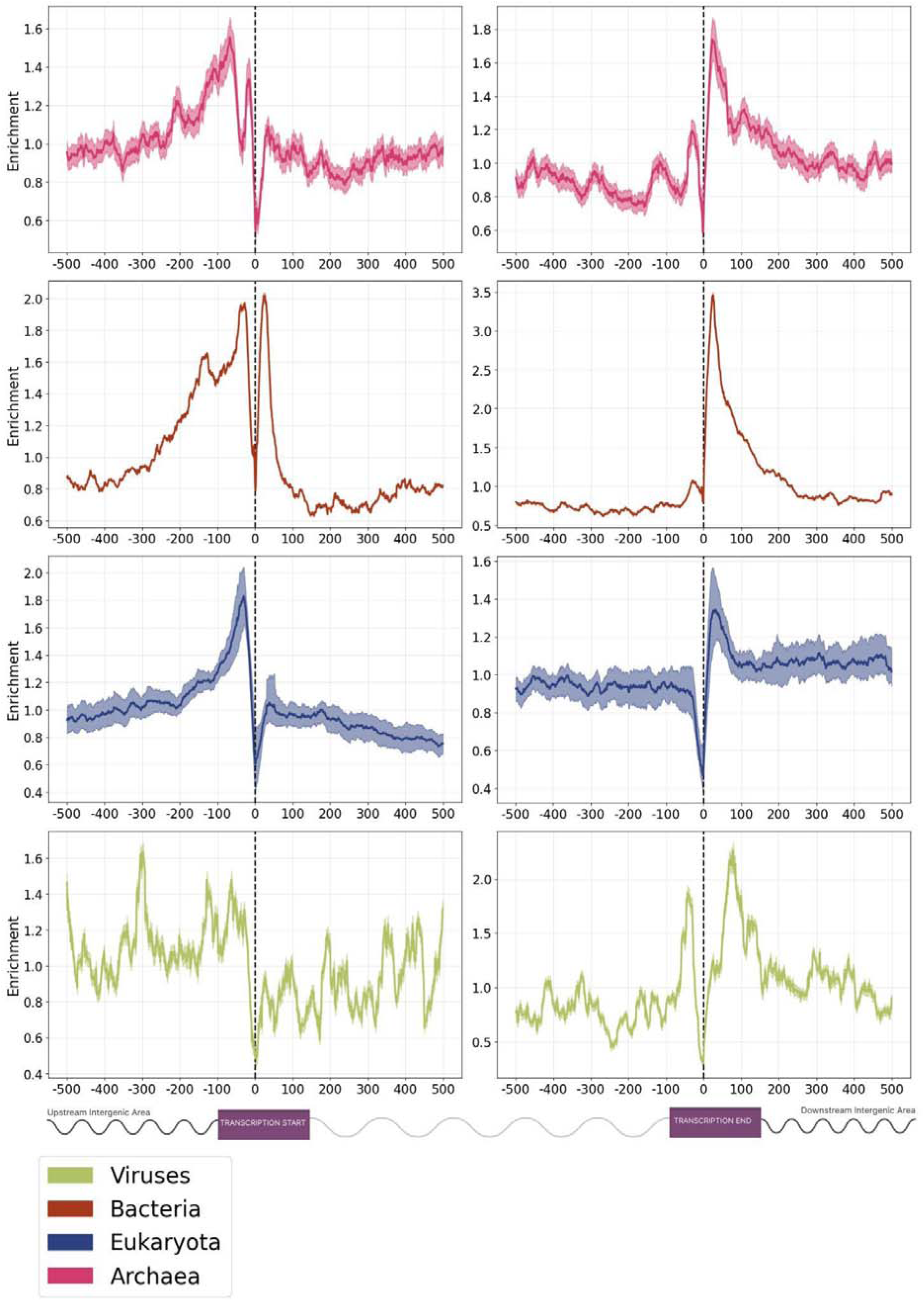
The topography of G4s relative to transcription start and transcription end sites across the tree of life. G4 distribution across the three domains of life and viruses derived using the regular expression-based algorithm. Confidence intervals represent the 2.5% lowest and 97.5% highest percentile from Monte-Carlo simulations with replacement (N=1,000).

**Supplementary Figure 6:**
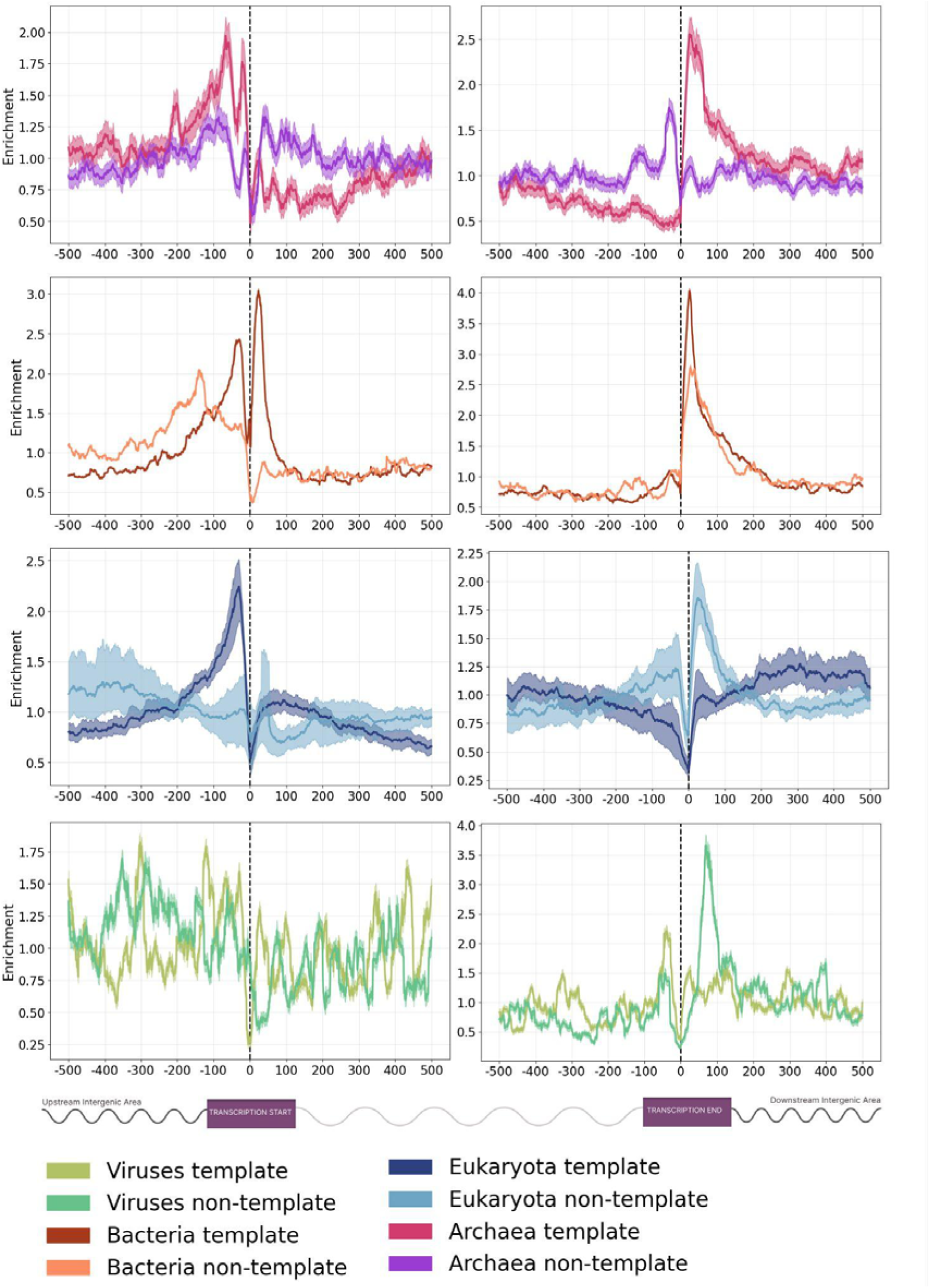
The topography of G4s relative to transcription start and transcription end sites across the tree of life at the template and non-template strands. G4 distribution across the three domains of life and viruses derived using the regular expression-based algorithm. Confidence intervals represent the 2.5% lowest and 97.5% highest percentile from Monte-Carlo simulations with replacement (N=1,000).

**Supplementary Figure 7:**
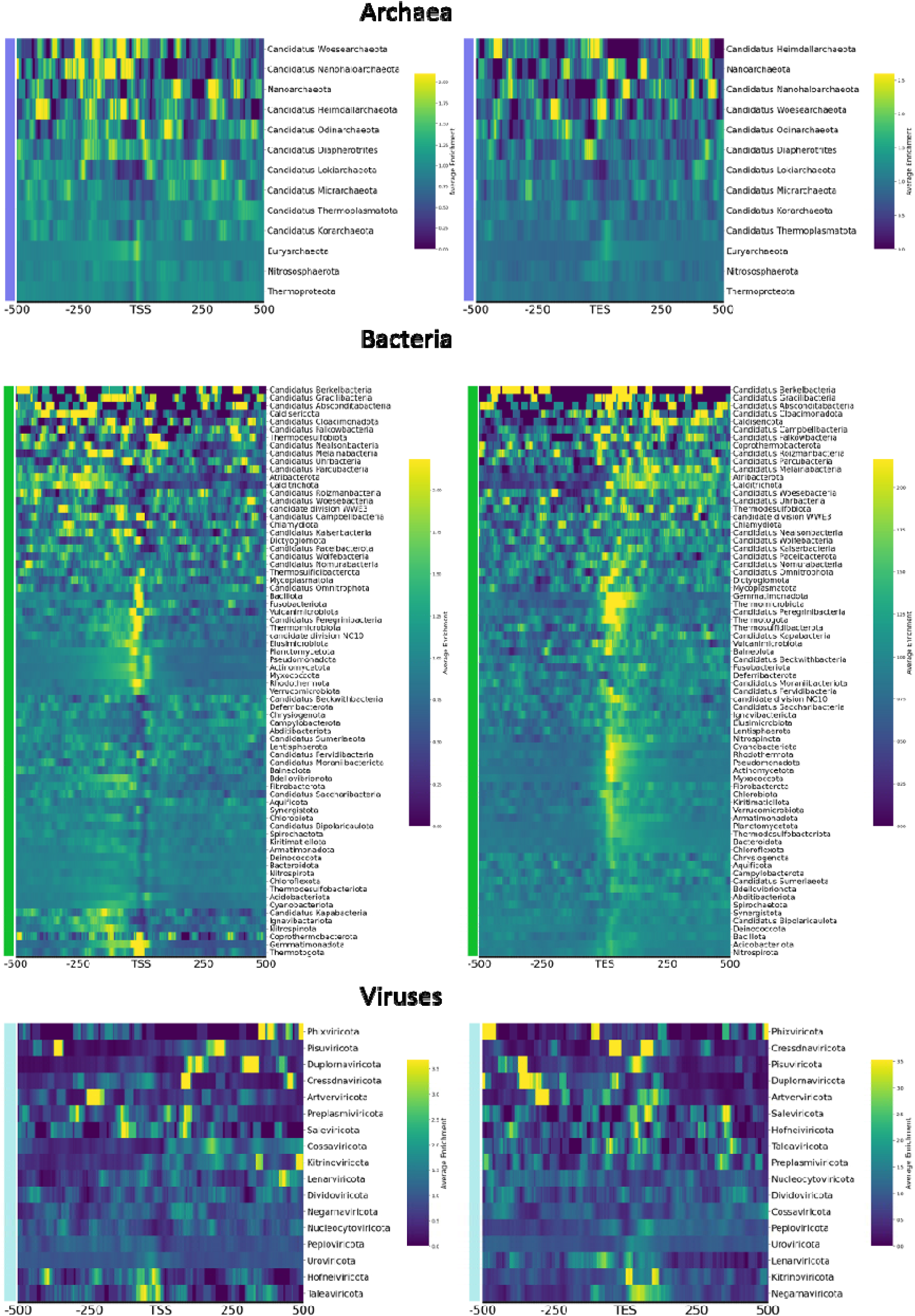
Distribution of G4s relative to transcription start and transcription end sites across archaeal, bacterial, and viral phyla. Results shown from the G4Hunter-based algorithm.

**Supplementary Figure 8:**
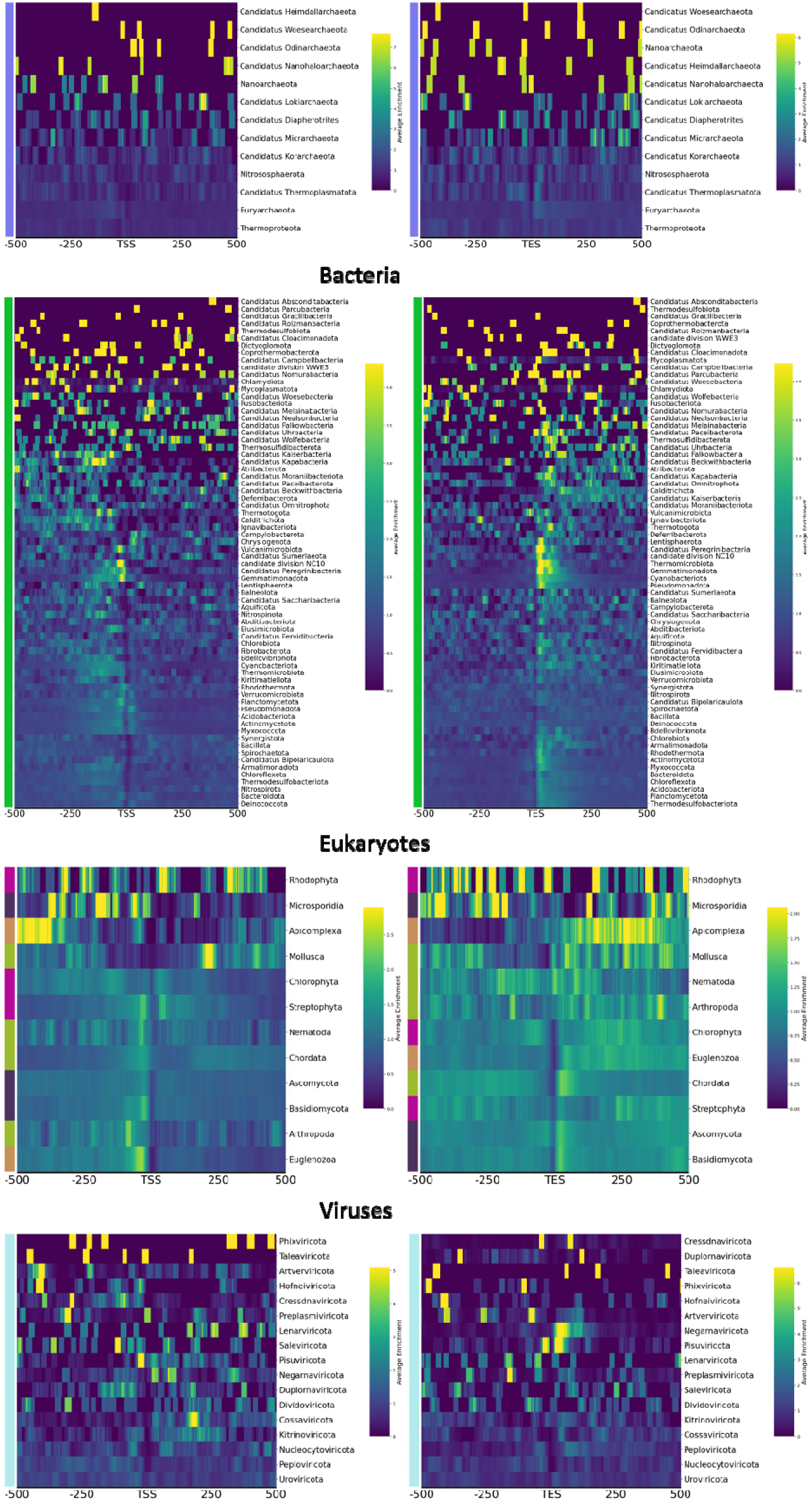
Distribution of G4s relative to transcription start and transcription end sites across archaeal, bacterial, and viral phyla. Results shown for the consensus motif from the regex-based algorithm.

**Supplementary Figure 9:**
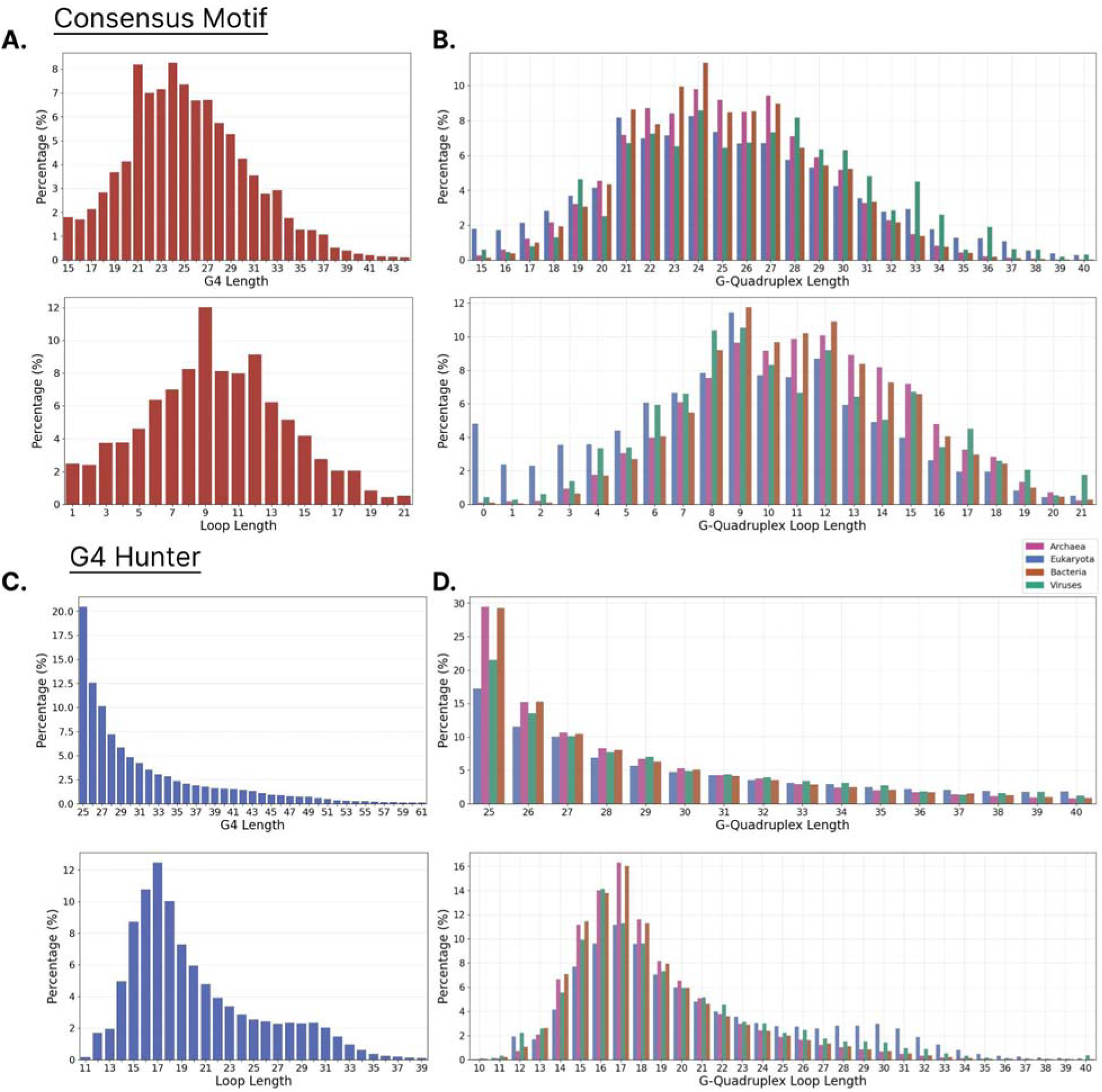
Length distribution of G4 lengths and of intervening loops of G4s across taxonomic groups. A-B) Distribution of G-quadruplex sequence length and of intervening loops for the consensus motif and from the consensus motif. **C-D)** Distribution of G-quadruplex total lengths and loop length for the three domains of life and viruses using the G4Hunter-based algorithm.

**Supplementary Figure 10:**
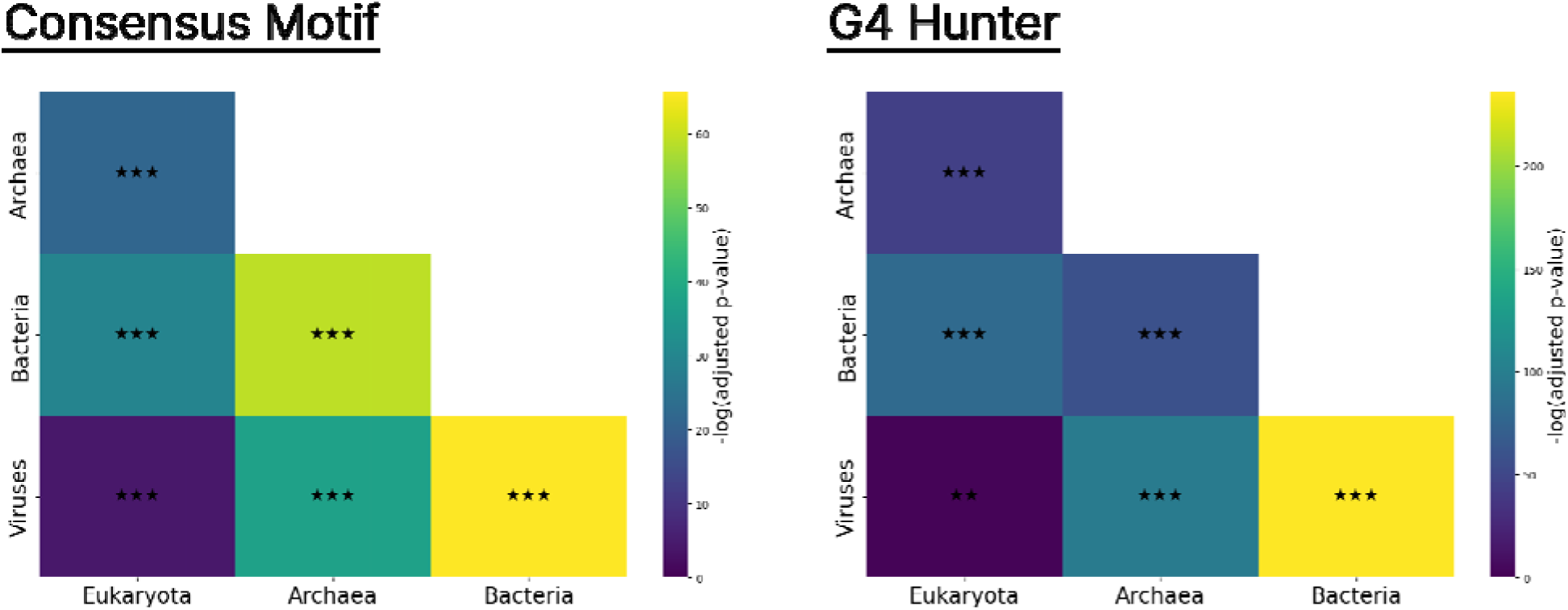
Biophysical properties of G4s in regards to loop length between taxonomic groups. Results show from the regex-based (consensus motif) and G4hunter-based algorithms.

**Supplementary Figure 11:**
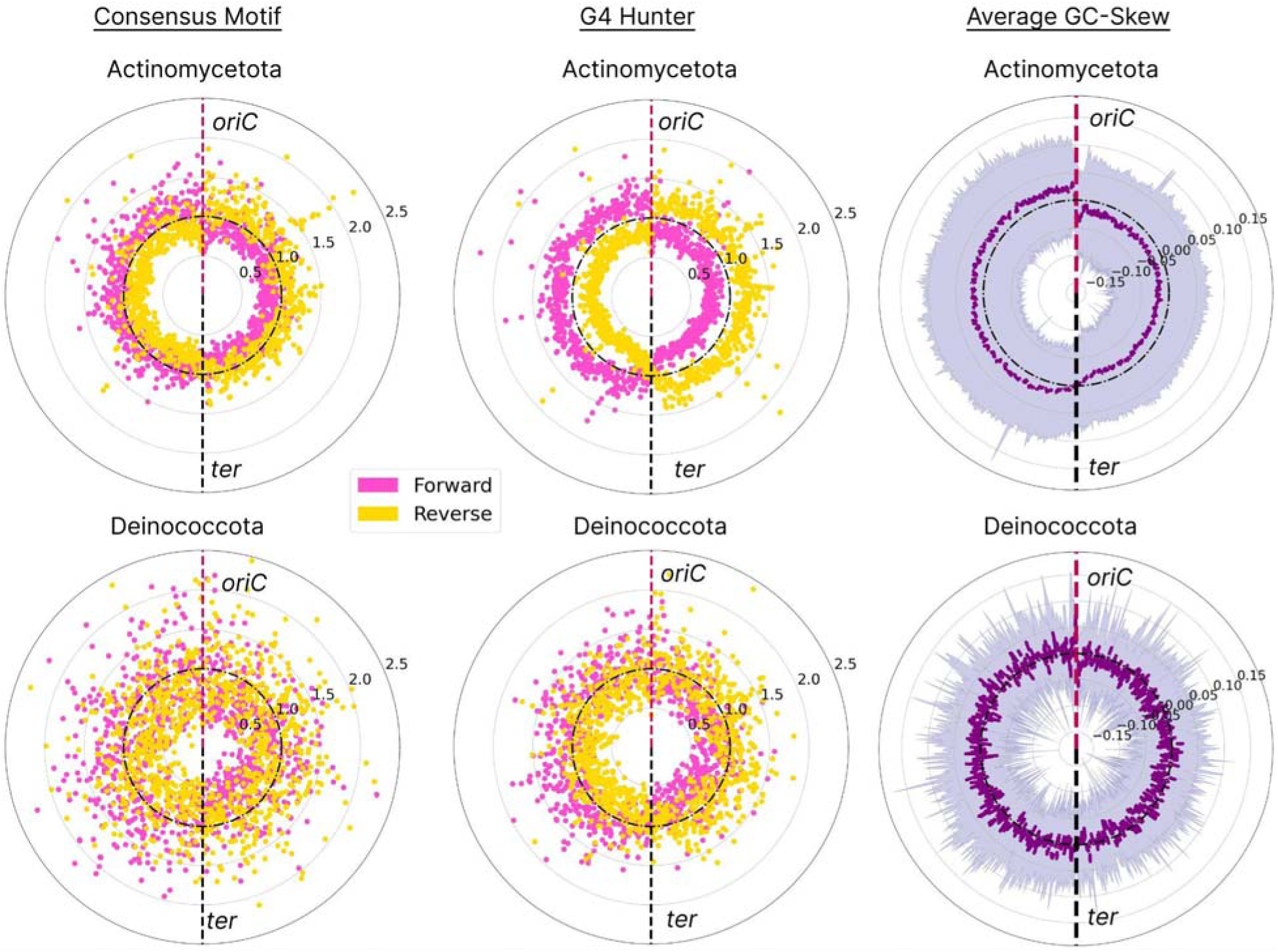
G4 distribution patterns relative to replication origin in bacterial phyla. Results shown for Actionomycetota and Deinococcota. G4s in forward and reverse strand orientation are shown in yellow and pink, respectively. Results shown for G4 Hunter-based and Consensus Motif-based algorithms. Average GC skew is also calculated and shown with purple.

**Supplementary Table 1:** Phyla and host of the top 100 viral genomes having the highest G4 densities.

**A.**
| Phylum | No. of Species | Total no. of G4s | Average G4 Density (per Mb) | Hosts |
| --- | --- | --- | --- | --- |
| Peploviricota | 73 | 38836 | 3489.00 | human, vertebrates |
| Kitrinoviricota | 11 | 384 | 4905.59 | fungi, land plants |
| Pisuviricota | 8 | 244 | 3731.96 | human, vertebrates |
| Artverviricota | 5 | 154 | 3463.75 | human, vertebrates |
| Dividoviricota | 1 | 63 | 3698.05 |  |
| Duplornaviricota | 1 | 20 | 3801.56 | vertebrates, invertebrates |
| Preplasmiviricota | 1 | 1 | 3676.47 |  |

**B.**
| Phylum | No. of Species | Total no. of G4s | Average G4 Density (per Mb) | Hosts |
| --- | --- | --- | --- | --- |
| Peploviricota | 94 | 38836 | 3489.00 | human, vertebrates |
| Kitrinoviricota | 4 | 384 | 4905.59 | land plants |
| Cressdnaviricota | 2 | 244 | 3731.96 | eukaryotic algae, invertebrates, vertebrates |

**Supplementary Table 2.**
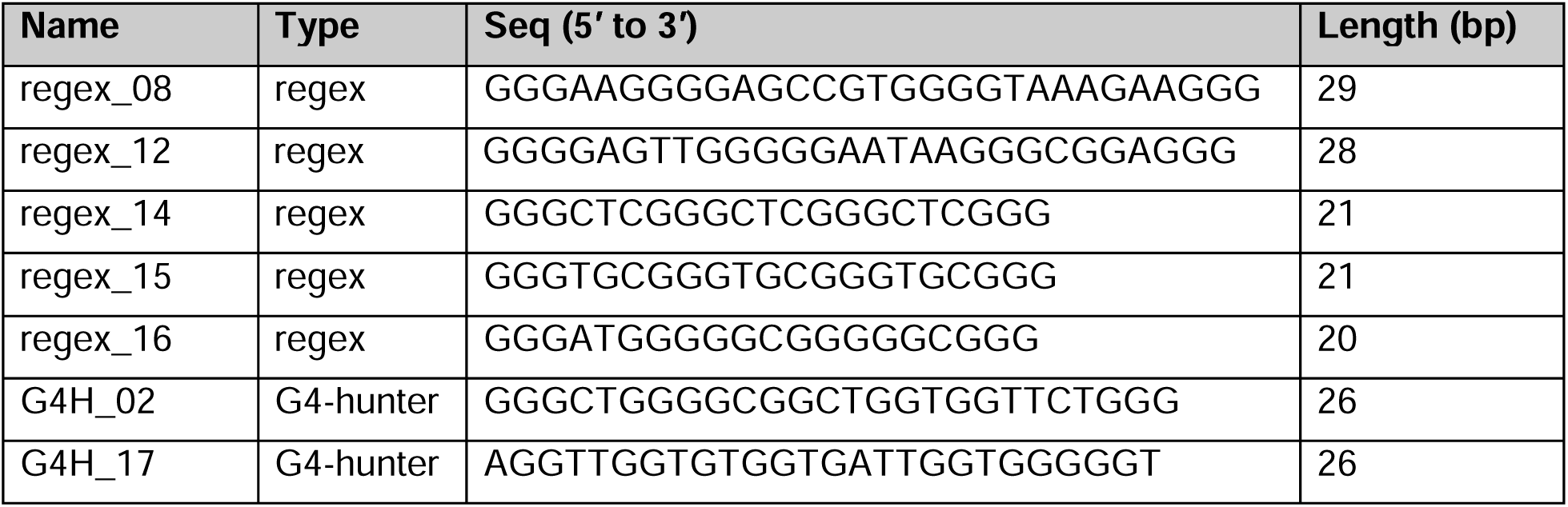
Selected sequences from the G4 database.

## References

1. Spiegel, J., Adhikari, S. and Balasubramanian, S. (2020) The Structure and Function of DNA G-Quadruplexes. Trends Chem, 2, 123–136.

2. Parkinson, G.N., Lee, M.P.H. and Neidle, S. (2002) Crystal structure of parallel quadruplexes from human telomeric DNA. Nature, 417, 876–880.

3. Spiegel, J., Cuesta, S.M., Adhikari, S., Hänsel-Hertsch, R., Tannahill, D. and Balasubramanian, S. (2021) G-quadruplexes are transcription factor binding hubs in human chromatin. Genome Biol., 22, 117.

4. Georgakopoulos-Soares, I., Victorino, J., Parada, G.E., Agarwal, V., Zhao, J., Wong, H.Y., Umar, M.I., Elor, O., Muhwezi, A., An, J.-Y., et al. (2022) High-throughput characterization of the role of non-B DNA motifs on promoter function. Cell Genom, 2.

5. Georgakopoulos-Soares, I., Chan, C.S.Y., Ahituv, N. and Hemberg, M. (2022) High-throughput techniques enable advances in the roles of DNA and RNA secondary structures in transcriptional and post-transcriptional gene regulation. Genome Biol., 23, 159.

6. Shen, J., Varshney, D., Simeone, A., Zhang, X., Adhikari, S., Tannahill, D. and Balasubramanian, S. (2021) Promoter G-quadruplex folding precedes transcription and is controlled by chromatin. Genome Biol., 22, 143.

7. Lago, S., Nadai, M., Cernilogar, F.M., Kazerani, M., Domíniguez Moreno, H., Schotta, G. and Richter, S.N. (2021) Promoter G-quadruplexes and transcription factors cooperate to shape the cell type-specific transcriptome. Nat. Commun., 12, 3885.

8. Brooks, T.A., Kendrick, S. and Hurley, L. (2010) Making sense of G-quadruplex and i-motif functions in oncogene promoters. FEBS J., 277, 3459–3469.

9. Hänsel-Hertsch, R., Beraldi, D., Lensing, S.V., Marsico, G., Zyner, K., Parry, A., Di Antonio, M., Pike, J., Kimura, H., Narita, M., et al. (2016) G-quadruplex structures mark human regulatory chromatin. Nat. Genet., 48, 1267–1272.

10. Li, G., Su, G., Wang, Y., Wang, W., Shi, J., Li, D. and Sui, G. (2023) Integrative genomic analyses of promoter G-quadruplexes reveal their selective constraint and association with gene activation. Commun Biol, 6, 625.

11. Robinson, J., Raguseo, F., Nuccio, S.P., Liano, D. and Di Antonio, M. (2021) DNA G-quadruplex structures: more than simple roadblocks to transcription? Nucleic Acids Res., 49, 8419–8431.

12. Huppert, J.L., Bugaut, A., Kumari, S. and Balasubramanian, S. (2008) G-quadruplexes: the beginning and end of UTRs. Nucleic Acids Res., 36, 6260–6268.

13. Lee, D.S.M., Ghanem, L.R. and Barash, Y. (2020) Integrative analysis reveals RNA G-quadruplexes in UTRs are selectively constrained and enriched for functional associations. Nat. Commun., 11, 527.

14. Georgakopoulos-Soares, I., Parada, G.E., Wong, H.Y., Medhi, R., Furlan, G., Munita, R., Miska, E.A., Kwok, C.K. and Hemberg, M. (2022) Alternative splicing modulation by G-quadruplexes. Nat. Commun., 13, 1–16.

15. Huang, H., Zhang, J., Harvey, S.E., Hu, X. and Cheng, C. (2017) RNA G-quadruplex secondary structure promotes alternative splicing via the RNA-binding protein hnRNPF. Genes Dev., 31, 2296–2309.

16. Ghosh, A., Pandey, S.P., Joshi, D.C., Rana, P., Ansari, A.H., Sundar, J.S., Singh, P., Khan, Y., Ekka, M.K., Chakraborty, D., et al. (2023) Identification of G-quadruplex structures in MALAT1 lncRNA that interact with nucleolin and nucleophosmin. Nucleic Acids Res., 51, 9415–9431.

17. Yuan, J., He, X. and Wang, Y. (2023) G-quadruplex DNA contributes to RNA polymerase II-mediated 3D chromatin architecture. Nucleic Acids Res., 51, 8434–8446.

18. Li, L., Williams, P., Ren, W., Wang, M.Y., Gao, Z., Miao, W., Huang, M., Song, J. and Wang, Y. (2021) YY1 interacts with guanine quadruplexes to regulate DNA looping and gene expression. Nat. Chem. Biol., 17, 161–168.

19. Wulfridge, P., Yan, Q., Rell, N., Doherty, J., Jacobson, S., Offley, S., Deliard, S., Feng, K., Phillips-Cremins, J.E., Gardini, A., et al. (2023) G-quadruplexes associated with R-loops promote CTCF binding. Mol. Cell, 83, 3064–3079.e5.

20. Georgakopoulos-Soares, I., Parada, G.E. and Hemberg, M. (2022) Secondary structures in RNA synthesis, splicing and translation. Comput. Struct. Biotechnol. J., 20, 2871–2884.

21. Murat, P., Marsico, G., Herdy, B., Ghanbarian, A.T., Portella, G. and Balasubramanian, S. (2018) RNA G-quadruplexes at upstream open reading frames cause DHX36- and DHX9-dependent translation of human mRNAs. Genome Biol., 19, 229.

22. Bugaut, A. and Balasubramanian, S. (2012) 5’-UTR RNA G-quadruplexes: translation regulation and targeting. Nucleic Acids Res., 40, 4727–4741.

23. Song, J., Perreault, J.-P., Topisirovic, I. and Richard, S. (2016) RNA G-quadruplexes and their potential regulatory roles in translation. Translation (Austin), 4, e1244031.

24. Kumari, S., Bugaut, A., Huppert, J.L. and Balasubramanian, S. (2007) An RNA G-quadruplex in the 5’ UTR of the NRAS proto-oncogene modulates translation. Nat. Chem. Biol., 3, 218– 221.

25. Lyu, K., Chow, E.Y.-C., Mou, X., Chan, T.-F. and Kwok, C.K. (2021) RNA G-quadruplexes (rG4s): genomics and biological functions. Nucleic Acids Res., 49, 5426–5450.

26. Georgakopoulos-Soares, I., Morganella, S., Jain, N., Hemberg, M. and Nik-Zainal, S. (2018) Noncanonical secondary structures arising from non-B DNA motifs are determinants of mutagenesis. Genome Res., 28, 1264–1271.

27. Wang, E., Thombre, R., Shah, Y., Latanich, R. and Wang, J. (2021) G-Quadruplexes as pathogenic drivers in neurodegenerative disorders. Nucleic Acids Res., 49, 4816–4830.

28. Wang, G. and Vasquez, K.M. (2023) Dynamic alternative DNA structures in biology and disease. Nat. Rev. Genet., 24, 211–234.

29. Simone, R., Fratta, P., Neidle, S., Parkinson, G.N. and Isaacs, A.M. (2015) G-quadruplexes: Emerging roles in neurodegenerative diseases and the non-coding transcriptome. FEBS Lett., 589, 1653–1668.

30. Zhang, R., Shu, H., Wang, Y., Tao, T., Tu, J., Wang, C., Mergny, J.-L. and Sun, X. (2023) G-Quadruplex Structures Are Key Modulators of Somatic Structural Variants in Cancers. Cancer Res., 83, 1234–1248.

31. Makova, K.D. and Weissensteiner, M.H. (2023) Noncanonical DNA structures are drivers of genome evolution. Trends Genet., 39, 109–124.

32. Kwok, C.K. and Merrick, C.J. (2017) G-Quadruplexes: Prediction, Characterization, and Biological Application. Trends Biotechnol., 35, 997–1013.

33. Ida, R. and Wu, G. (2008) Direct NMR detection of alkali metal ions bound to G-quadruplex DNA. J. Am. Chem. Soc., 130, 3590–3602.

34. Harkness, R.W.,5th and Mittermaier, A.K. (2017) G-quadruplex dynamics. *Biochim. Biophys*. Acta: Proteins Proteomics, 1865, 1544–1554.

35. Chan, C.-Y., Umar, M.I. and Kwok, C.K. (2019) Spectroscopic analysis reveals the effect of a single nucleotide bulge on G-quadruplex structures. Chem. Commun., 55, 2616–2619.

36. Del Villar-Guerra, R., Trent, J.O. and Chaires, J.B. (2018) G-Quadruplex Secondary Structure Obtained from Circular Dichroism Spectroscopy. Angew. Chem. Int. Ed Engl., 57, 7171– 7175.

37. Mergny, J.-L. and Lacroix, L. (2009) UV Melting of G-Quadruplexes. *Curr. Protoc. Nucleic Acid Chem.*, Chapter 17, 17.1.1–17.1.15.

38. Chambers, V.S., Marsico, G., Boutell, J.M., Di Antonio, M., Smith, G.P. and Balasubramanian, S. (2015) High-throughput sequencing of DNA G-quadruplex structures in the human genome. Nat. Biotechnol., 33, 877–881.

39. Marsico, G., Chambers, V.S., Sahakyan, A.B., McCauley, P., Boutell, J.M., Antonio, M.D. and Balasubramanian, S. (2019) Whole genome experimental maps of DNA G-quadruplexes in multiple species. Nucleic Acids Res., 47, 3862–3874.

40. Kwok, C.K., Marsico, G., Sahakyan, A.B., Chambers, V.S. and Balasubramanian, S. (2016) rG4-seq reveals widespread formation of G-quadruplex structures in the human transcriptome. Nat. Methods, 13, 841–844.

41. Zhao, J., Chow, E.Y.-C., Yeung, P.Y., Zhang, Q.C., Chan, T.-F. and Kwok, C.K. (2022) Enhanced transcriptome-wide RNA G-quadruplex sequencing for low RNA input samples with rG4-seq 2.0. BMC Biol., 20, 257.

42. Di Antonio, M., Ponjavic, A., Radzevičius, A., Ranasinghe, R.T., Catalano, M., Zhang, X., Shen, J., Needham, L.-M., Lee, S.F., Klenerman, D., et al. (2020) Single-molecule visualization of DNA G-quadruplex formation in live cells. Nat. Chem., 12, 832–837.

43. Summers, P.A., Lewis, B.W., Gonzalez-Garcia, J., Porreca, R.M., Lim, A.H.M., Cadinu, P., Martin-Pintado, N., Mann, D.J., Edel, J.B., Vannier, J.B., et al. (2021) Visualising G-quadruplex DNA dynamics in live cells by fluorescence lifetime imaging microscopy. Nat. Commun., 12, 162.

44. Huppert, J.L. and Balasubramanian, S. (2005) Prevalence of quadruplexes in the human genome. Nucleic Acids Res., 33, 2908–2916.

45. Lombardi, E.P. and Londoño-Vallejo, A. (2020) A guide to computational methods for G-quadruplex prediction. Nucleic Acids Res., 48, 1603.

46. Bedrat, A., Lacroix, L. and Mergny, J.-L. (2016) Re-evaluation of G-quadruplex propensity with G4Hunter. Nucleic Acids Res., 44, 1746–1759.

47. Sahakyan, A.B., Chambers, V.S., Marsico, G., Santner, T., Di Antonio, M. and Balasubramanian, S. (2017) Machine learning model for sequence-driven DNA G-quadruplex formation. Sci. Rep., 7, 14535.

48. Bartas, M., Čutová, M., Brázda, V., Kaura, P., Šťastný, J., Kolomazník, J., Coufal, J., Goswami, P., Červeň, J. and Pečinka, P. (2019) The Presence and Localization of G-Quadruplex Forming Sequences in the Domain of Bacteria. Molecules, 24.

49. Rawal, P., Kummarasetti, V.B.R., Ravindran, J., Kumar, N., Halder, K., Sharma, R., Mukerji, M., Das, S.K. and Chowdhury, S. (2006) Genome-wide prediction of G4 DNA as regulatory motifs: role in Escherichia coli global regulation. Genome Res., 16, 644–655.

50. Brázda, V., Luo, Y., Bartas, M., Kaura, P., Porubiaková, O., Šťastný, J., Pečinka, P., Verga, D., Da Cunha, V., Takahashi, T.S., et al. (2020) G-Quadruplexes in the Archaea Domain. Biomolecules, 10.

51. Saranathan, N. and Vivekanandan, P. (2019) G-Quadruplexes: More Than Just a Kink in Microbial Genomes. Trends Microbiol., 27, 148–163.

52. Métifiot, M., Amrane, S., Litvak, S. and Andreola, M.-L. (2014) G-quadruplexes in viruses: function and potential therapeutic applications. Nucleic Acids Res., 42, 12352–12366.

53. Lavezzo, E., Berselli, M., Frasson, I., Perrone, R., Palù, G., Brazzale, A.R., Richter, S.N. and Toppo, S. (2018) G-quadruplex forming sequences in the genome of all known human viruses: A comprehensive guide. PLoS Comput. Biol., 14, e1006675.

54. Zhong, H.-S., Dong, M.-J. and Gao, F. (2023) G4Bank: A database of experimentally identified DNA G-quadruplex sequences. Interdiscip. Sci., 15, 515–523.

55. Vannutelli, A., Schell, L.L.N., Perreault, J.-P. and Ouangraoua, A. (2023) GAIA: G-quadruplexes in alive creature database. Nucleic Acids Res., 51, D135–D140.

56. Yu, H., Qi, Y., Yang, B., Yang, X. and Ding, Y. (2023) G4Atlas: a comprehensive transcriptome-wide G-quadruplex database. Nucleic Acids Res., 51, D126–D134.

57. Qian, S.H., Shi, M.-W., Xiong, Y.-L., Zhang, Y., Zhang, Z.-H., Song, X.-M., Deng, X.-Y. and Chen, Z.-X. (2023) EndoQuad: a comprehensive genome-wide experimentally validated endogenous G-quadruplex database. Nucleic Acids Res., 10.1093/nar/gkad966.

58. Ghosh, A., Largy, E. and Gabelica, V. (2021) DNA G-quadruplexes for native mass spectrometry in potassium: a database of validated structures in electrospray-compatible conditions. Nucleic Acids Res., 49, 2333–2345.

59. Wang, Y.-H., Yang, Q.-F., Lin, X., Chen, D., Wang, Z.-Y., Chen, B., Han, H.-Y., Chen, H.-D., Cai, K.-C., Li, Q., et al. (2022) G4LDB 2.2: a database for discovering and studying G-quadruplex and i-Motif ligands. Nucleic Acids Res., 50, D150–D160.

60. Li, Q., Xiang, J.-F., Yang, Q.-F., Sun, H.-X., Guan, A.-J. and Tang, Y.-L. (2013) G4LDB: a database for discovering and studying G-quadruplex ligands. Nucleic Acids Res., 41, D1115–23.

61. Zok, T., Kraszewska, N., Miskiewicz, J., Pielacinska, P., Zurkowski, M. and Szachniuk, M. (2022) ONQUADRO: a database of experimentally determined quadruplex structures. Nucleic Acids Res., 50, D253–D258.

62. O’Leary, N.A., Wright, M.W., Brister, J.R., Ciufo, S., Haddad, D., McVeigh, R., Rajput, B., Robbertse, B., Smith-White, B., Ako-Adjei, D., et al. (2016) Reference sequence (RefSeq) database at NCBI: current status, taxonomic expansion, and functional annotation. Nucleic Acids Res., 44, D733–45.

63. Benson, D.A., Cavanaugh, M., Clark, K., Karsch-Mizrachi, I., Lipman, D.J., Ostell, J. and Sayers, E.W. (2013) GenBank. Nucleic Acids Res., 41, D36–42.

64. Dong, M.-J., Luo, H. and Gao, F. (2023) DoriC 12.0: an updated database of replication origins in both complete and draft prokaryotic genomes. Nucleic Acids Res., 51, D117– D120.

65. Huppert, J.L. and Balasubramanian, S. (2007) G-quadruplexes in promoters throughout the human genome. Nucleic Acids Res., 35, 406–413.

66. Brázda, V., Kolomazník, J., Lýsek, J., Bartas, M., Fojta, M., Šťastný, J. and Mergny, J.-L. (2019) G4Hunter web application: a web server for G-quadruplex prediction. Bioinformatics, 35, 3493–3495.

67. Steinegger, M. and Söding, J. (2018) Clustering huge protein sequence sets in linear time. Nat. Commun., 9, 1–8.

68. Steinegger, M. and Söding, J. (2017) MMseqs2 enables sensitive protein sequence searching for the analysis of massive data sets. Nat. Biotechnol., 35, 1026–1028.

69. Katoh, K., Misawa, K., Kuma, K. and Miyata, T. (2002) MAFFT: a novel method for rapid multiple sequence alignment based on fast Fourier transform. Nucleic Acids Res., 30, 3059–3066.

70. Eddy, S.R. (2011) Accelerated Profile HMM Searches. PLoS Comput. Biol., 7, e1002195.

71. Berman, H.M., Westbrook, J., Feng, Z., Gilliland, G., Bhat, T.N., Weissig, H., Shindyalov, I.N. and Bourne, P.E. (2000) The Protein Data Bank. Nucleic Acids Res., 28, 235–242.

72. Camacho, C., Coulouris, G., Avagyan, V., Ma, N., Papadopoulos, J., Bealer, K. and Madden, T.L. (2009) BLAST+: architecture and applications. BMC Bioinformatics, 10, 1–9.

73. Yachdav, G., Wilzbach, S., Rauscher, B., Sheridan, R., Sillitoe, I., Procter, J., Lewis, S.E., Rost, B. and Goldberg, T. (2016) MSAViewer: interactive JavaScript visualization of multiple sequence alignments. Bioinformatics, 32, 3501–3503.

74. Wheeler, T.J., Clements, J. and Finn, R.D. (2014) Skylign: a tool for creating informative, interactive logos representing sequence alignments and profile hidden Markov models. BMC Bioinformatics, 15, 1–9.

75. Sehnal, D., Bittrich, S., Deshpande, M., Svobodová, R., Berka, K., Bazgier, V., Velankar, S., Burley, S.K., Koča, J. and Rose, A.S. (2021) Mol* Viewer: modern web app for 3D visualization and analysis of large biomolecular structures. Nucleic Acids Res., 49, W431– W437.

76. Wheeler, T.J. and Eddy, S.R. (2013) nhmmer: DNA homology search with profile HMMs. Bioinformatics, 29, 2487–2489.

77. Li, Z., Qian, S.H., Wang, F., Mohamed, H.I., Yang, G., Chen, Z.-X. and Wei, D. (2022) G-quadruplexes in genomes of viruses infecting eukaryotes or prokaryotes are under different selection pressures from hosts. J. Genet. Genomics, 49, 20–29.

78. Chen, M.C., Tippana, R., Demeshkina, N.A., Murat, P., Balasubramanian, S., Myong, S. and Ferré-D’Amaré, A.R. (2018) Structural basis of G-quadruplex unfolding by the DEAH/RHA helicase DHX36. Nature, 558, 465–469.

79. Guiblet, W.M., DeGiorgio, M., Cheng, X., Chiaromonte, F., Eckert, K.A., Huang, Y.-F. and Makova, K.D. (2021) Selection and thermostability suggest G-quadruplexes are novel functional elements of the human genome. Genome Res., 31, 1136–1149.

80. Grigoriev, A. (1998) Analyzing genomes with cumulative skew diagrams. Nucleic Acids Res., 26, 2286–2290.

81. Merrikh, C.N. and Merrikh, H. (2018) Gene inversion potentiates bacterial evolvability and virulence. Nat. Commun., 9, 4662.

82. Kota, S., Dhamodharan, V., Pradeepkumar, P.I. and Misra, H.S. (2015) G-quadruplex forming structural motifs in the genome of Deinococcus radiodurans and their regulatory roles in promoter functions. Appl. Microbiol. Biotechnol., 99, 9761–9769.

83. Frasson, I., Soldà, P., Nadai, M., Tassinari, M., Scalabrin, M., Gokhale, V., Hurley, L.H. and Richter, S.N. (2022) Quindoline-derivatives display potent G-quadruplex-mediated antiviral activity against herpes simplex virus 1. Antiviral Res., 208, 105432.

84. Warner, E.F., Bohálová, N., Brázda, V., Waller, Z.A.E. and Bidula, S. (2021) Analysis of putative quadruplex-forming sequences in fungal genomes: novel antifungal targets? Microb Genom, 7.

85. Watanabe, S. (2020) Cyanobacterial multi-copy chromosomes and their replication. Biosci. Biotechnol. Biochem., 84, 1309–1321.

86. Ohbayashi, R., Hirooka, S., Onuma, R., Kanesaki, Y., Hirose, Y., Kobayashi, Y., Fujiwara, T., Furusawa, C. and Miyagishima, S.-Y. (2020) Evolutionary Changes in DnaA-Dependent Chromosomal Replication in Cyanobacteria. Front. Microbiol., 11, 786.

87. Estep, K.N., Butler, T.J., Ding, J. and Brosh, R.M. (2019) G4-Interacting DNA Helicases and Polymerases: Potential Therapeutic Targets. Curr. Med. Chem., 26, 2881–2897.

88. Puig Lombardi, E., Holmes, A., Verga, D., Teulade-Fichou, M.-P., Nicolas, A. and Londoño-Vallejo, A. (2019) Thermodynamically stable and genetically unstable G-quadruplexes are depleted in genomes across species. Nucleic Acids Res., 47, 6098–6113.

89. Guiblet, W.M., Cremona, M.A., Harris, R.S., Chen, D., Eckert, K.A., Chiaromonte, F., Huang, Y.-F. and Makova, K.D. (2021) Non-B DNA: a major contributor to small- and large-scale variation in nucleotide substitution frequencies across the genome. Nucleic Acids Res., 49, 1497–1516.

90. Wu, F., Niu, K., Cui, Y., Li, C., Lyu, M., Ren, Y., Chen, Y., Deng, H., Huang, L., Zheng, S., et al. (2021) Genome-wide analysis of DNA G-quadruplex motifs across 37 species provides insights into G4 evolution. Commun Biol, 4, 98.

91. Puig Lombardi, Emilia, Arturo Londoño-Vallejo, and Alain Nicolas. 2019. “Relationship Between G-Quadruplex Sequence Composition in Viruses and Their Hosts.” Molecules 24 (10). 10.3390/molecules24101942.

